# A conserved cytoplasmic glutamic acid mediates pH-dependent gating in bacterial urea channel UreI

**DOI:** 10.64898/2025.12.08.692916

**Authors:** Anna Stoib, Sahar Shojaei, Xenia Fischer, Sandra Posch, Gianluca Parisse, Mario Frezzini, Tobias Putz, Christine Siligan, Nikolaus Goessweiner-Mohr, Daniele Narzi, Andreas Horner

## Abstract

The bacterial UreI channel family enables rapid urea uptake, essential for urease activity and acid resistance. While pH-dependent gating in *Helicobacter pylori* UreI (*Hp*UreI) is attributed to periplasmic histidines, the role of cytoplasmic residues remains unexplored. The UreI homolog from *Streptococcus salivarius* (*Ss*UreI), lacking periplasmic histidines, serves as a simplified model to identify gating determinants. Here, we combine yeast complementation assays, in vitro studies, and MD simulations to show that cytoplasmic glutamic acid E136 mediates pH-dependent gating in *Ss*UreI. Mutagenesis in *H. pylori*/*H. hepaticus* homologs confirm E136 as a conserved pH sensor across UreI channels. Protonation of E136 disrupts its salt bridge with R20, increasing cytoplasmic-loop flexibility and urea-permeable filter conformations. These findings challenge the paradigm of exclusive periplasmic pH sensing, supporting a Gram-negative-specific dual-sensor model (E136 + histidines). By elucidating this molecular mechanism, we identify E136 as a therapeutic target to disrupt UreI-mediated acid resistance in pathogenic bacteria.

---

Urea serves as a critical nitrogen source for bacteria and plays a central role in survival, physiology, and adaptation to environmental stresses, including acidic conditions. In pathogenic ureolytic bacteria such as *Helicobacter pylori*, urea uptake and metabolism are essential for colonization and survival in the acidic environment of the stomach. Chronic *H. pylori* infection, which affects ∼50% of the global population, is a major risk factor for gastric ulcers and stomach cancer^1^. In the oral cavity, dynamic pH fluctuations arise from bacterial metabolism of dietary carbohydrates, favoring the growth of aciduric pathogens like *Streptococcus mutans*, a primary contributor to dental caries^2^. Conversely, ureolytic bacteria such as *Streptococcus salivarius* mitigate acidification by hydrolyzing urea into ammonia, thereby neutralizing the local environment and suppressing acidogenic species^3^. While this process reduces caries risk^3^, the resulting alkaline shift can also promote mineral deposition and calculus formation, contributing to gingivitis and periodontitis^4,5^.

*S. salivarius*, a commensal organism prevalent on oral soft tissues, employs the UreI/urease system to counteract glycolytic acidification. UreI facilitates urea uptake into the cytoplasm, where urease converts it to ammonia and CO₂, buffering both intracellular and extracellular pH^6^. Complementary mechanisms, including a plasma membrane-bound ATPase (proton expulsion) and cyclopropane fatty acid synthase (reduced membrane proton permeability), further support pH homeostasis^3^. However, the relative contributions of these mechanisms, particularly the energy allocation for urea hydrolysis versus proton expulsion, remain unresolved.

The UreI channel family is central to urea uptake across bacterial species. *Hp*UreI, the pH-gated urea channel of *H. pylori*, enables survival in the acidic gastric environment (pH ∼2) by facilitating urea uptake for hydrolysis into ammonia and CO₂, thereby buffering the periplasmic space^7,8^. *Hp*UreI exhibits pH-dependent gating, opening at pH 5 and closing at pH 7 (pKₐ ∼5.9), ensuring urease activity is restricted to acidic conditions and preventing excessive cytoplasmic alkalization^8,9^. The prevailing hypothesis implicates periplasmic histidine residues as the pH sensor in *Hp*UreI^10,11^. However, this model fails to explain gating in homologs lacking periplasmic histidines, such as UreI from *S. salivarius* (*Ss*UreI) (**Figure 1**), which shares significant sequence homology with *Hp*UreI but was previously reported to function pH-independently in vitro^12^. This discrepancy is puzzling, given *Ss*UreI’s essential role in modulating intracellular and extracellular pH in acidic environments, where pH-dependent regulation would be critical for survival.

**Figure 1.**
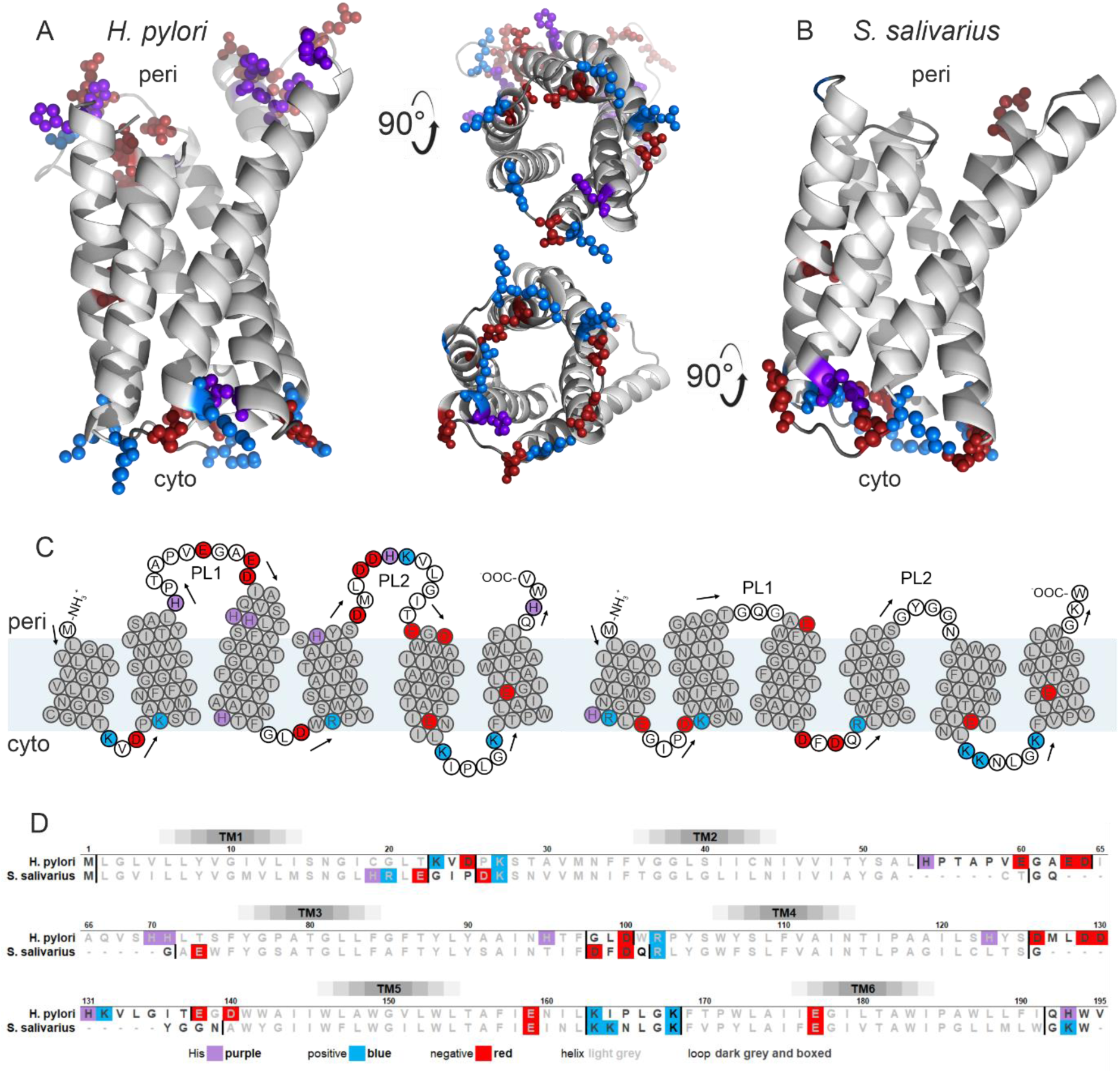
Structural and sequence comparison of SsUreI and HpUreI. **A.** Alphafold 3 structure of HpUreI. HpUreI (195 amino acids, ∼21 kDa) folds into six transmembrane helices (TM1–6), with its N- and C-termini and two large periplasmic loops (PL1 and PL2) oriented toward the periplasm. Since PL1 is unresolved in X-ray^8^ and cryo-EM^11^ structures, an AlphaFold3 model^13^ is shown. Protonatable residues (His: purple; Arg/Lys: blue; Glu/Asp: red) are highlighted on both the periplasmic and cytoplasmic sides. **B.** Structure of SsUreI. An AlphaFold3 model of the SsUreI protomer (171 amino acids, ∼19 kDa) is shown in side view (periplasmic entrance facing upward) and top view (cytoplasmic side). Unlike HpUreI, SsUreI lacks large periplasmic loops and periplasmic protonatable residues. Protonatable residues are represented as spheres, color-coded as in (A). **C.** Topology comparison. Snake plot representation of HpUreI and SsUreI, illustrating their transmembrane helix organization and loop regions according to the Alphafold 3 structures. **D.** Sequence alignment. Alignment of SsUreI and HpUreI sequences, with protonatable residues highlighted to emphasize conserved and divergent features.

To resolve this paradox, we employed *Ss*UreI, a simplified UreI homolog lacking periplasmic histidines, as a model system to dissect the molecular basis of pH gating. Using a multidisciplinary approach, combining yeast complementation assays, stopped-flow measurements with reconstituted *Ss*UreI, all-atom MD simulations, and comparative sequence/structural analyses, we demonstrate that *Ss*UreI is indeed pH-gated, with a cytoplasmic glutamic acid residue (E136) acting as the primary pH sensor. Comparative experiments with homologous channels from *H. hepaticus* and *H. pylori* suggest that E136 plays a conserved role in bacterial proton-gated urea channels. These results challenge the prevailing hypothesis of exclusive periplasmic pH sensing and provide a molecular framework for understanding acid adaptation in both commensal and pathogenic bacteria. By identifying E136 as the pH sensor, this study elucidates the gating mechanism of UreI channels and opens avenues for targeting them in pathogenic bacteria, where they are critical for survival and virulence.

## Results

### SsUreI exhibits pH-dependent gating, challenging prior assumptions of pH independence

Previous studies reported that *Ss*UreI facilitates passive urea transport independently of ambient pH (pH 4.0–8.0)^12^. Structurally, *Ss*UreI differs from its homolog *Hp*UreI by featuring short periplasmic loops and harboring only one protonatable periplasmic residue (E59) between transmembrane helices TM2 and TM3. Like *Hp*UreI, *Ss*UreI contains a glutamic acid residue (E154) within its selectivity filter. Based on these structural differences, periplasmic histidine residues were proposed to mediate pH gating in *Hp*UreI^8,10,11^. However, initial yeast complementation assays provided the first evidence that this assumption might be incorrect, as *Ss*UreI exhibited pH-dependent urea transport akin to *Hp*UreI.

To investigate this discrepancy, we expressed *Ss*UreI in the urea uptake-deficient yeast strain *S. cerevisiae* YNVW1 (Δdur3), where growth in liquid media depends on urea uptake via the exogenous transporter (**Figure 2A**). Under these conditions, urea serves as the sole nitrogen source, and cell growth reflects the functional activity of *Ss*UreI. As expected, yeast expressing *Ss*UreI exhibited a sigmoidal growth pattern: robust growth at acidic pH (high urea permeability) and minimal growth at neutral pH (low urea permeability) (**Figure 2B**). Normalizing these data to cell concentrations in arginine-containing media^9^ revealed a pH-dependent gating profile with a pK_a_ of 5.7 ± 0.06 (**Figure 2C**). Contrary to prior reports^12^, these results demonstrate that *Ss*UreI is a pH-gated urea channel, similar to *Hp*UreI.

**Figure 2.**
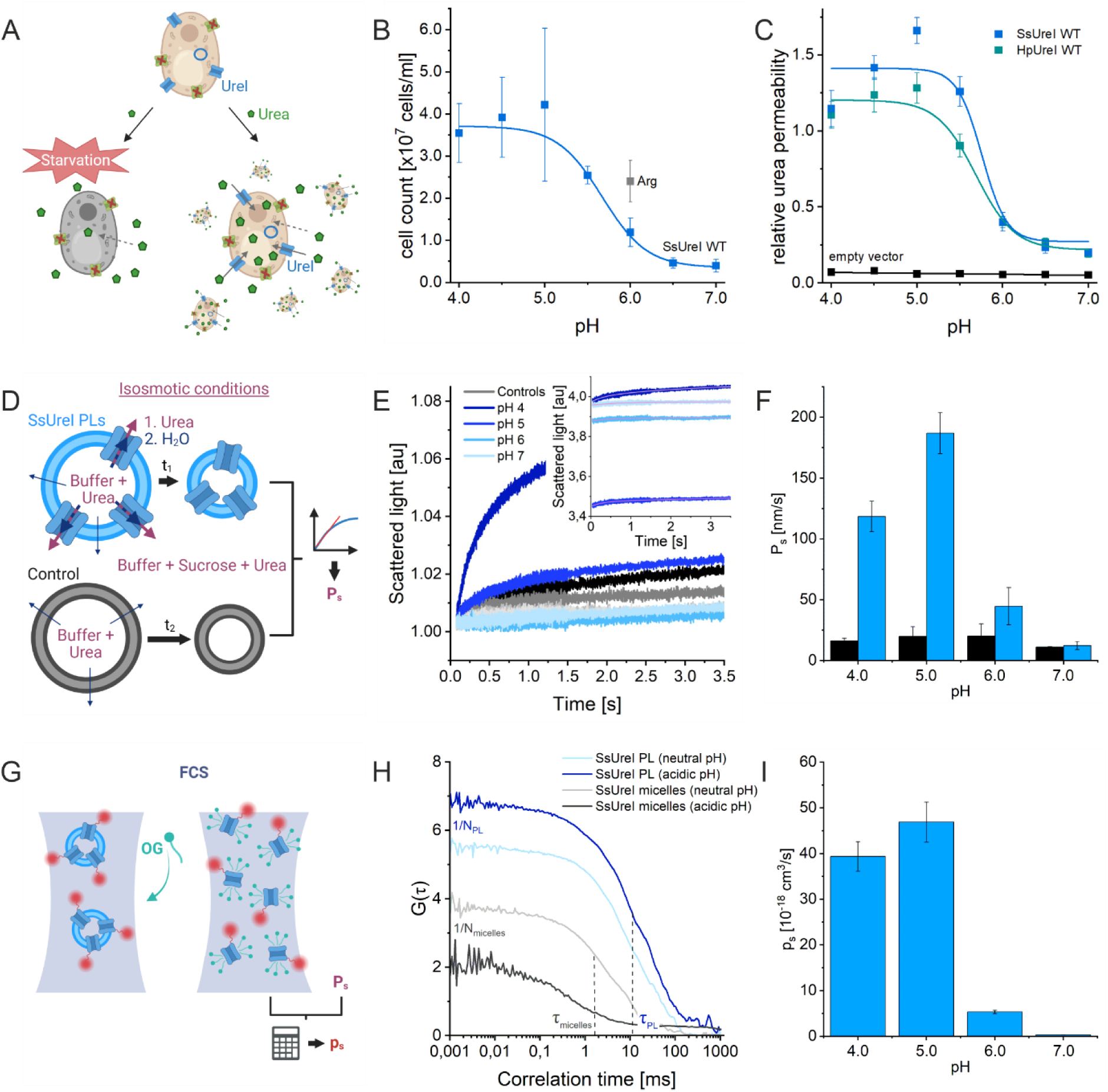
SsUreI is a pH-gated urea channel. **A.** Yeast growth assay for urea permeability. The urea uptake-deficient strain S. cerevisiae Δdur3 expressing SsUreI can grow only if the channel facilitates urea transport. Higher urea permeability results in increased cell growth, while low passive membrane permeability (dotted arrows) leads to starvation. **B.** Cell counts of S. cerevisiae Δdur3 expressing SsUreI WT, measured after 2 days of incubation in urea media at pH 4.0–7.0 or arginine media at pH 6.0 (internal standard, gray). An exemplary dataset for SsUreI WT (average cell count ± standard deviation (SD)) in urea (blue) and arginine (grey) media is shown. **C**. Normalized urea permeabilities (± SEM) for SsUreI WT (n=6), HpUreI WT (n=18), and empty vector (n=7, negative control). Data demonstrate pH-dependent gating of SsUreI, with maximal permeability at acidic pH. **D.** Schematic of the stopped-flow vesicle shrinkage assay. A urea gradient drives urea efflux from vesicles through reconstituted SsUreI, while sucrose (impermeable) remains outside, maintaining osmolarity. **E.** Scattered light signal increases as vesicles shrink due to water efflux following urea permeation. The signal change reflects vesicle volume reduction and refractive index changes^19^. **F.** Averaged urea permeabilities (± SD) for SsUreI-containing LUVs (PLUVs) and bare lipid vesicles (controls) across three independent purification/reconstitution experiments. Data confirm pH-dependent urea transport through SsUreI. **G.** FCS quantifies SsUreI protomers per vesicle by comparing AF647-labeled SsUreI particles before/after OG (octylglucoside) solubilization. **H.** Exemplary FCS autocorrelation curves for SsUreI PLUVs at neutral and acidic pH. OG addition increases particle number (N) (lower autocorrelation amplitude) and reduces diffusion time (τ) indicating vesicle disruption. **I.** pH-dependent unitary urea permeability *p*_*s*_ of SsUreI. The trend mirrors the pH dependence observed in yeast growth assays (C), confirming pH-gated urea transport.

### In vitro assays confirm pH-dependent gating of SsUreI

To validate these findings, we purified and reconstituted *Ss*UreI into large unilamellar vesicles (LUVs) and measured its pH-dependent unitary urea permeability using a combination of stopped-flow spectroscopy, dynamic light scattering (DLS), and fluorescence correlation spectroscopy (FCS). In this assay, urea efflux from vesicles under isosmotic conditions drives osmotic water loss, causing vesicle shrinkage (**Figure 2D**). The rate of shrinkage, monitored via scattered light intensity, directly correlates with urea permeability across the vesicle membrane.

*Ss*UreI-containing vesicles exhibited higher urea permeability at acidic pH (pH 5.0) compared to neutral pH (pH 7.0), as evidenced by faster light-scattering changes (**Figure 2E**). Fitting these data to an exponential model (**Eqn. 2**) yielded membrane urea permeabilities (*P*_*s*_) that varied with pH, forming a bell-shaped curve with maximal permeability at pH 5.0, reduced permeability at pH 4.0, and minimal permeability at pH 7.0 (**Figure 2E**). This trend mirrors the pH-dependent growth observed in yeast, confirming that *Ss*UreI’s gating is pH-regulated. Control experiments in yeast verified that the *Ss*UreI purification construct (*Ss*UreIcl) exhibited gating behavior identical to wild-type *Ss*UreI (**Extended Data Figure 1**). Finally, we used FCS^14-19^ to quantify the number of *Ss*UreI hexamers per vesicle (**Figure 2G** and **H**) and calculated the unitary urea permeability (*p*_*s*_) using **Eqn. 3**. The maximum unitary permeability in the open state was determined to be 47 ± 6 × 10⁻¹⁸ cm³/s (**Figure 2I**).

### SsUreI facilitates pH-dependent ammonia transport, mirroring its urea permeability properties

Previous studies have demonstrated that *Hp*UreI not only mediates pH-dependent urea transport (pK_a_ of 5.53 ± 0.003) but also pH-dependent ammonia permeability^9^. This dual permeability may enable *H. pylori* to neutralize periplasmic acidity by facilitating the back-diffusion of ammonia, a hydrolysis product of cytoplasmic urease, into the periplasm. To investigate whether *Ss*UreI shares this property, we expressed it in the ammonia uptake-deficient yeast strain *S. cerevisiae* Sc18-Δmep1-3 (lacking endogenous ammonia transporters MEP1-3). As with our urea complementation assays, we monitored yeast growth in liquid media at varying pH levels, with ammonia as the sole nitrogen source (**Extended Data Figure 2A**).

If *Ss*UreI facilitates ammonia transport, yeast cells proliferate; otherwise, they starve due to insufficient ammonia uptake. *Ss*UreI-expressing yeast exhibited a pH-dependent growth pattern, with maximal growth and ammonia permeability at pH 5.5 and minimal growth at neutral pH (**Extended Data Figure 2B**). Normalizing these data to cell concentrations in arginine-containing media^9^ revealed a pH-dependent gating profile for ammonia, with a pK_a_ of 5.8 ± 0.16 (**Extended Data Figure 2C**). These results mirror those observed for urea transport and confirm that *Ss*UreI, like *Hp*UreI, is a pH-gated ammonia channel. The pH-dependent ammonia permeability of *Ss*UreI suggests a dual role in acid resistance: facilitating urea uptake for cytoplasmic ureolysis and ammonia back-diffusion to neutralize oral acidity. However, due to the high intrinsic passive permeability of biological membranes to ammonia^20,21^, we were unable to reproduce these findings *in vitro*.

### SsUreI exhibits negligible water permeability across a physiological pH range

Many membrane channels and transporters, including aquaporins, facilitate passive water transport^16,22,23^. To assess whether *Ss*UreI shares this property, we examined its water permeability using a freeze-survival assay (**Extended Data Figure 2D**) in yeast lacking endogenous water channels (strain Y20000). In this assay, water-permeable channels (e.g., aquaporins) enhance cell survival by mitigating osmotic stress during freezing as less ice crystals are formed in the cells^24,25^. Survival was monitored using FDA (live cells, green fluorescence) and PI (dead cells, red fluorescence). *Ss*UreI-expressing yeast showed negligible survival, comparable to empty vector controls and contrasting with hAQP1-expressing cells (positive control; **Extended Data Figure 2E**). While *Hp*UreI showed a slight increase in survival, it remained significantly lower than *h*AQP1 (**Extended Data Figure 2F**), suggesting minimal water permeability for *Ss*UreI.

To validate these findings, we subjected *Ss*UreI-containing LUVs, the same batch used for urea permeability assays, to an osmotic gradient (sucrose addition). Water efflux from vesicles, driven by the osmotic gradient, was monitored via light scattering (**Extended Data Figure 2G-H)**. As shown in **Extended Data Figure 2I**, *Ss*UreI-containing vesicles exhibited negligible water permeability across the pH range of 4.0–7.0, comparable to empty vesicles. For context, vesicles containing a single AQP1 tetramer exhibit a *P*_*f*_ > 30 µm/s under similar conditions^15^. Thus, we conclude that *Ss*UreI does not facilitate significant water transport. The lack of water permeability in *Ss*UreI suggests a unique role in acid resistance in *S. salivarius*.

### Cytoplasmic E136 and R20 mediate pH-dependent gating in SsUreI

pH-dependent gating in membrane channels is typically driven by protonation or deprotonation of amino acid side chains. Given the pK_a_ of *Ss*UreI channel opening/closing (∼5.7), we focused on residues with side-chain pK_a_ values near this range: histidine (H, pK_a_ 6.04), aspartic acid (D, pK_a_ 3.7), and glutamic acid (E, pK_a_ 4.2). The local protein environment can shift these pK_a_ values, potentially aligning them with the channel’s gating pK_a_. To identify the pH-sensing residues, we mutated all negatively charged residues and cytoplasmic histidines to their neutral, sterically similar counterparts (H→F, E→Q, D→N) and assessed their pH-dependent urea permeability using yeast complementation assays (**Figure 3A**).

**Figure 3.**
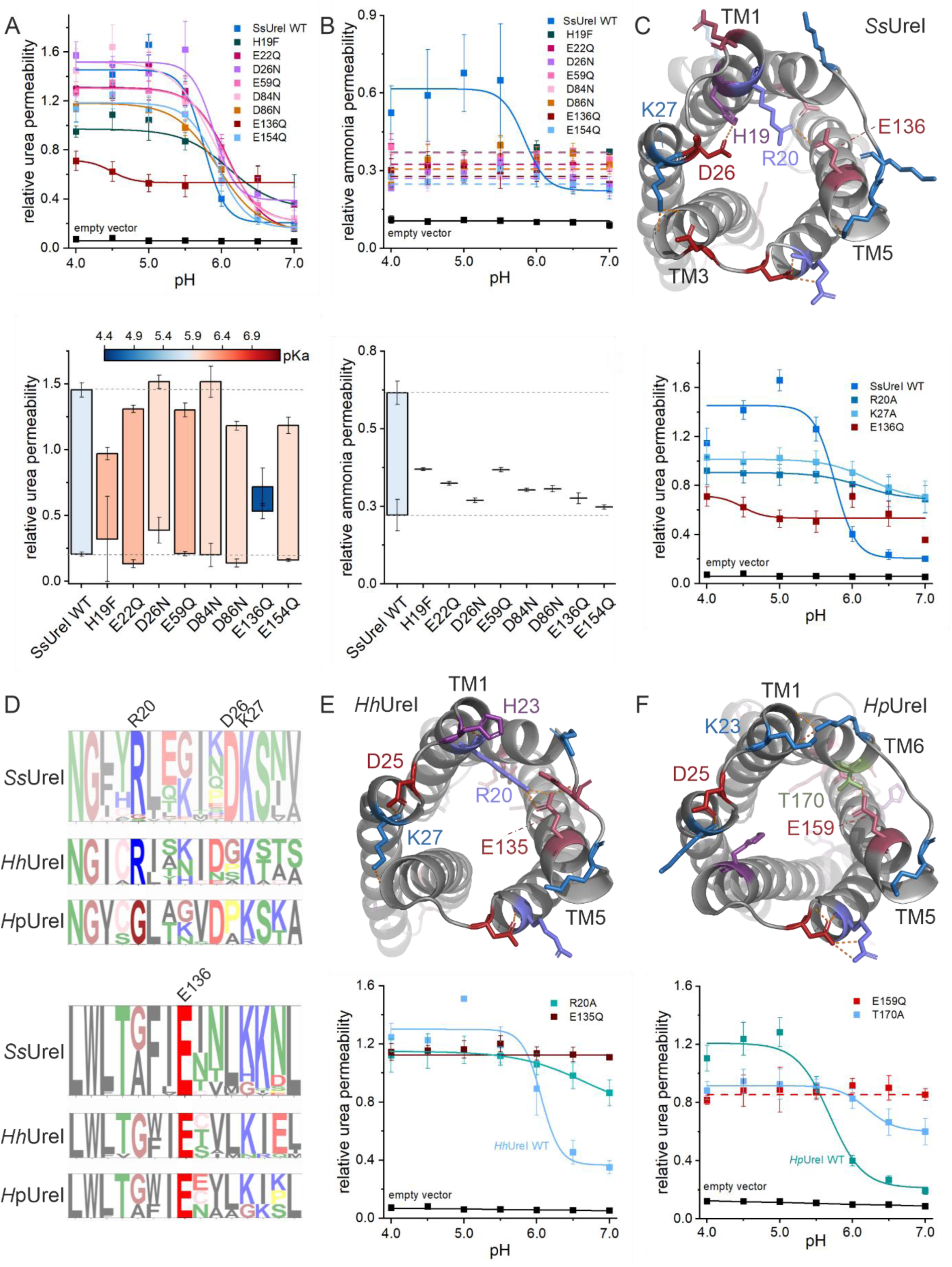
Cytoplasmic E136 constitutes the pH sensor of SsUreIs. **A.** pH-dependent urea permeability of SsUreI variants. Protonatable residues His (H), Glu (E), and Asp (D) were mutated to their neutral counterparts Phe (F), Gln (Q), and Asn (N), respectively. Relative urea permeabilities of SsUreI variants reveal shifts in pH-dependent gating (bars) and pK_a_ of channel opening/closing (color-coded) (pK_a_ values are listed in **Extended Data Table 1**). E136Q nearly abolishes pH gating, while H19F has a minor effect. **B.** pH-dependent ammonia permeability of SsUreI variants. All SsUreI mutants exhibit pH-independent, reduced ammonia permeability compared to wild-type SsUreI, suggesting that ammonia transport requires a precisely aligned selectivity filter. **C.** Cytoplasmic snapshot of SsUreI (AlphaFold3 model). Potential salt bridges (R20-E136, H19-D26) and H-bonds (K27-T80/I81) are highlighted. Mutations R20A and K27A disrupt gating similarly to E136Q, confirming their role in pH sensing and structural stability. **D.** Conservation of pH-sensing residues across UreI homologs. Logo plots of cytoplasmic loops 1 (CL1 between TM1–TM2) and 3 (CL3 between TM5–TM6) show that R20 and E136 (SsUreI) are highly conserved in enterohepatic Helicobacter UreI channels (e.g., HhUreI). In gastric Helicobacter (e.g., HpUreI), E136 is strictly conserved, while R20 is substituted by T170 in HpUreI (**Extended Data** Figures 3-5). **E.** Impaired gating in HhUreI mutants. Mutations R20A and E135Q (homologous to E136Q in SsUreI) in HhUreI disrupt pH gating, mirroring the effects observed in SsUreI. **F.** Impaired gating in HpUreI mutants. Mutations E159Q (homologous to E136Q) and T170A (interaction partner of E159) in HpUreI impair pH gating, analogous to HhUreI and SsUreI.

While absolute permeability comparisons between open and closed states were not feasible due to potential expression level variations, two mutations, H19F and E136Q, significantly altered *Ss*UreI’s pH-dependent activity. E136Q rendered the channel largely pH-independent, whereas H19F subtly disrupted gating. In contrast, ammonia permeability in these mutants differed markedly from urea permeability (**Figure 3B**). Any mutation converted *Ss*UreI into a pH-independent weak ammonia channel, suggesting that ammonia transport requires a precisely aligned selectivity filter, less tolerant to structural perturbations than urea transport. This aligns with observations in *Hp*UreI^9^ and implies that urea and ammonia may traverse the channel via distinct pathways.

### Structural analysis reveals a conserved cytoplasmic gating mechanism

Inspection of the AlphaFold3-predicted *Ss*UreI structure (**Figure 3C**) suggested that E136 (TM5) and H19 (TM1) participate in salt-bridge formation with neighboring residues. We hypothesized that E136 interacts with R20 and H19 with D26. To test this, we mutated the proposed binding partners (D26N and R20A) and assessed their gating behavior. While D26N showed no gating alterations, R20A abolished pH-dependent gating, mirroring the effect of E136Q (**Figure 3C**). This confirms that E136 and R20 form a cytoplasmic pH-sensing pair in *Ss*UreI.

Additionally, K27 (TM1) forms a potential H-bond with the backbone of T80 and I81 (TM3). The K27A mutation disrupted pH gating similarly to R20A, suggesting that stabilizing interactions between TM1 and TM3 are critical for effective gating. Thus, pH-dependent gating in *Ss*UreI requires a structurally intact protomer, where E136-R20 interactions act as the primary cytoplasmic pH sensor.

### E136 is conserved and functionally critical across bacterial UreI channels

To determine whether E136’s role in pH gating extends to other bacterial UreI channels, we analyzed sequence conservation of R20 and E136 in homologs from Gram-positive and Gram-negative bacteria, including *H. hepaticus* (*Hh*UreI) and *H. pylori* (*Hp*UreI) (**Figure 3D**). E136 is perfectly conserved across all 68 UreI sequences analyzed, while R20 is conserved only in *Ss*UreI and *Hh*UreI homologs. Structural analysis of *Hh*UreI (AlphaFold3) revealed a similar E135-R20 interaction (**Figure 3E**). In *Hp*UreI, T170 substitutes for the missing R20, forming an H-bond with E159 (**Figure 3F**). This substitution suggests that structural plasticity in the cytoplasmic gating mechanism allows for functional conservation despite sequence variability.

To test the functional importance of these residues, we introduced R20A and E135Q mutations in *Hh*UreI and E159Q and T170A mutations in *Hp*UreI. Strikingly, all four mutations impaired pH-dependent gating, analogous to the effects observed in our model channel *Ss*UreI. This demonstrates that E136 (and its homologs E135/E159) acts as a conserved cytoplasmic pH sensor across bacterial UreI channels, sensing proton concentrations at the cytoplasmic entrance. Despite being a novel finding for UreI channels, cytoplasmic pH-sensing mechanisms are already observed in other channel families, including aquaporins^26-28^ and potassium channels^29-31^.

### Atomistic Simulations Reveal the Structural Basis of pH-Dependent Gating in SsUreI

To elucidate the molecular mechanism of pH-dependent gating in *Ss*UreI, we complemented our biochemical and mutagenesis data with atomistic molecular dynamics (MD) simulations of the *Ss*UreI hexamer embedded in a POPC bilayer (**Figure 4A–B**). Our goal was to dissect how protonation of key residues, particularly E136, modulates the channel’s conformational dynamics and its ability to permeate urea. We performed three ensembles of simulations, each consisting of four independent 1 µs trajectories initiated from distinct velocities, totaling 12 µs of sampling. This approach allowed us to robustly explore the conformational space under three protonation conditions: Standard protonation (SP) with all residues in their typical ionization states; Protonated E154 (P-E154) with E154 (selectivity filter) protonated; Protonated E154 and E136 (P-E154/E136) with both residues protonated (**Figure 4C**). All simulated systems remained structurally stable throughout the trajectories, as evidenced by backbone RMSD values < 0.3 nm for the hexamer and individual protomers (**Figures S4A**, **S5A**, **S6A**), consistent hydrogen-bond networks (**Figures S4B**, **S5B**, **S6B**), indicating preserved secondary and quaternary structure, a stable radius of gyration (**Figure S7**), and an intact central lipid plug occluding the hexameric pore as demonstrated by the time-resolved density profiles along the membrane normal (**Figure S8**).

**Figure 4.**
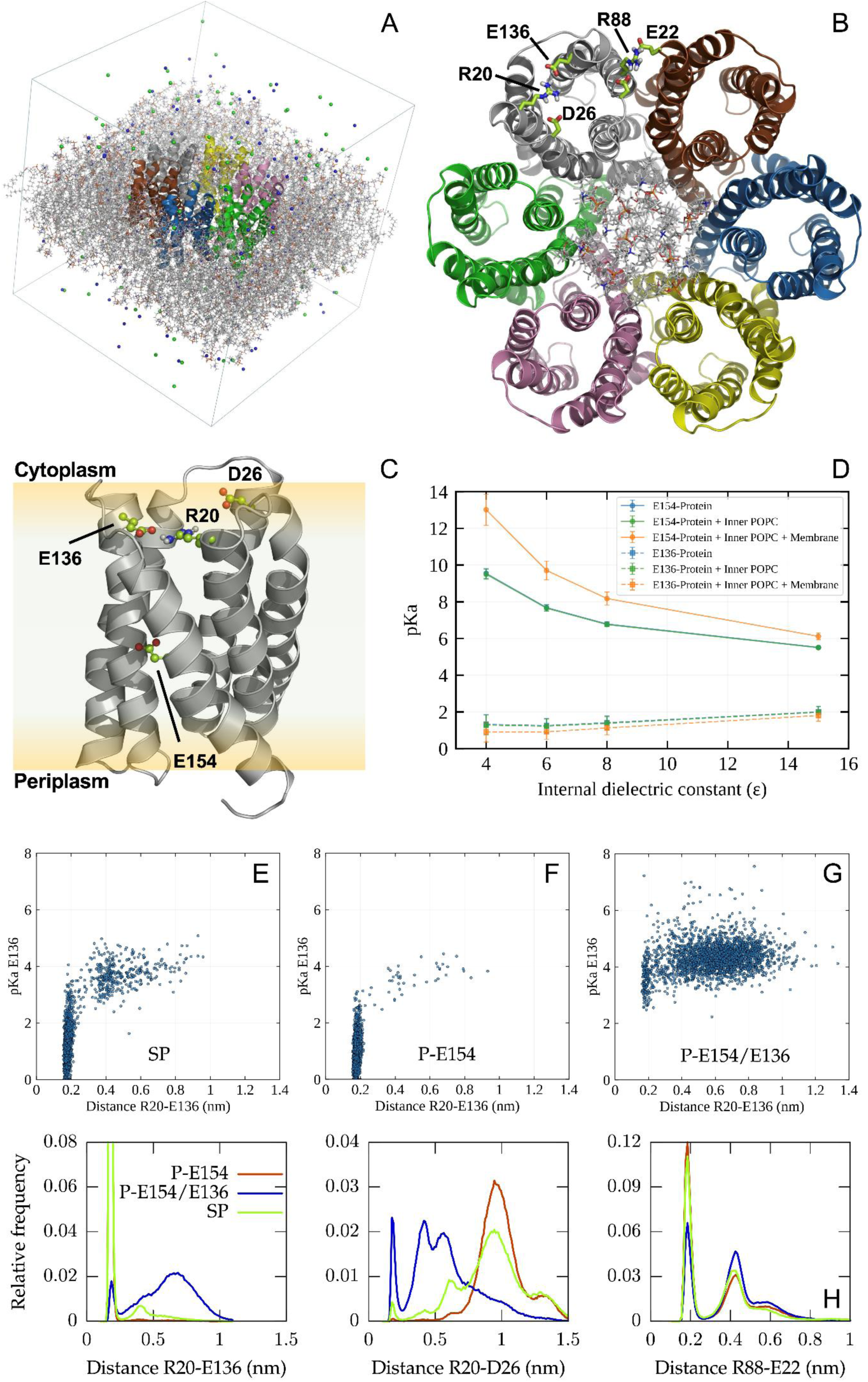
Atomistic simulations reveal coupling between protonation, residue interactions, and dynamics in SsUreI. **A.** Simulation system setup. The SsUreI hexamer (cartoon) is embedded in a POPC bilayer (sticks), with Na⁺/Cl⁻ ions (spheres) representing physiological ionic strength. **B.** Cytoplasmic view of the SsUreI hexamer. Six peripheral pores surround a central lipid plug (POPC lipids shown as sticks: nine cytoplasmic, three periplasmic). Protomers are individually coloured, and key residues (E136, R20, E154) are highlighted in sticks. **C.** Side view of a single protomer. Key residues (E136, R20, E154, D26) are shown in sticks, illustrating their spatial arrangement within the transmembrane region. **D.** pK_a_ sensitivity to dielectric environment. Predicted pK_a_ values for E154 and E136 in the starting model as a function of the dielectric constant used in Poisson–Boltzmann (PB) calculations. The pK_a_ of E154 decreases with higher dielectric constants, while E136 remains largely unaffected. **E–G.** Correlation between E136–R20 distance and pK_a_. pK_a_ values of E136, sampled every 10 ns during SP (E), P-E154 (F), and P-E154/E136 (G) simulations, are plotted against the minimum distance between the R20 guanidinium and E136 carboxylate groups. Each system includes 2,400 data points (100 frames × 4 replicas × 6 protomers), revealing how salt-bridge disruption elevates the pK_a_ of E136. **H.** Impact of E136 protonation on interaction networks. Minimum distance distributions between R20–E136 (left), R20–D26 (middle), and R88–E22 (right) highlight how E136 protonation destabilizes the E136–R20 salt bridge, promotes R20–D26 interactions, and weakens the inter-protomer R88–E22 interaction.

### Protonation of E136 and E154 Modulates pK_a_ and Conformational Dynamics

We first examined the system where all titratable residues were in their standard protonation states. To identify residues that might undergo protonation changes under physiological conditions, we calculated pK_a_ values for all titratable sites along the MD trajectories, sampling every 10 ns with a dielectric constant of 6. This yielded 2.400 pK_a_ estimates (100 frames × 4 replicas × 6 protomers) per residue. While tyrosine, arginine, and lysine residues consistently showed pK_a_ values > 9.0, E136 and E154 exhibited the greatest variability.

For E154, the only membrane-buried, pore-lining titratable residue (**Figure 4C**), the average predicted pK_a_ was ∼5.4 across trajectories. Protonation of E154 in the selectivity filter may alter its interactions with permeating solutes, potentially affecting the channel’s substrate specificity or gating kinetics. However, pK_a_ calculations on the starting structure revealed sensitivity to the dielectric constant and membrane environment (**Figure 4D**). With a dielectric constant of 6 (mimicking the protein interior), E154’s pK_a_ exceeded 7 in a protein-only environment and >9 when the membrane was explicitly included. Since Poisson–Boltzmann pK_a_ calculations typically approximate the membrane environment by adjusting the dielectric constant rather than modelling it explicitly, conventional estimates may underestimate the true proton affinity of the membrane-buried E154. Thus, E154 was considered protonated under both acidic (pH 5.0) and neutral/basic (pH 7.5) conditions in subsequent simulations. This assumption is supported by our yeast complementation assays, where the E154Q mutation didn’t show any impaired gating behavior, making a role of E154 protonation/deprotonation on *Ss*UreIs gating behavior unplausible.

In contrast, E136 exhibited a much lower average pK_a_ (∼1.6), though sporadic estimates reached ∼5. Intriguingly, these elevated pK_a_ values correlated with transient disruptions of the E136-R20 salt bridge. When the E136-R20 distance increased and the salt bridge broke, E136’s predicted pK_a_ rose (**Figure 4E**). This suggests that local conformational fluctuations render E136 protonatable within the physiological pH range (4.0–7.0). Based on these findings, we simulated two additional systems: P-E154 with E154 protonated and E136 deprotonated; P-E154/E136 with both E154 and E136 protonated.

### Protonation of E136 Disrupts Critical Interaction Networks

pK_a_ calculations on the trajectories of the P-E154 and P-E154/E136 systems revealed distinct protonation behaviors for the two key residues. E154 exhibited an average pK_a_ of 7.2–7.3, consistent with its role as a membrane-buried residue in the selectivity filter. E136 showed a mean pK_a_ of 1.0 in the P-E154 system (deprotonated) and 4.3 in the P-E154/E136 system (protonated). While these values are limited by the approximations inherent to continuum electrostatics, such as dielectric constant assumptions and fixed protonation states in classical MD, they provide qualitative insights into how local conformational states modulate pK_a_ values.

A central finding of this analysis is the critical role of the E136–R20 salt bridge in regulating E136’s protonation (**Figure 4E–G**). When the E136–R20 distance increased and the salt bridge was disrupted, the pK_a_ of E136 rose, suggesting that conformational dynamics directly influence its protonation state.

### E136 Protonation Alters Intra- and Inter-Protomer Interactions

Building on this observation, we examined how E136 protonation alters the broader interaction network at the cytosolic exit of the channel. In systems where E136 was deprotonated, the E136–R20 salt bridge remained stable (**Extended Data Table 2**). Protonation of E136 not only abolished the E136–R20 interaction but also allowed R20 to sample alternative conformations, including transient contacts with D26. Moreover, E136 protonation destabilized the inter-protomer salt bridge between R88 and E22 of adjacent protomers (**Figure 4H**). This suggests that the protonation state of E136 regulates not only local intra-protomer interactions but also quaternary contacts across the hexamer. These findings demonstrate that E136 acts as a conformational switch, coupling local protonation events to global rearrangements that open or close the cytoplasmic gate, thereby regulating urea permeation. This mechanism explains how physiological pH fluctuations (4.0–7.0) can modulate *Ss*UreI activity, enabling bacteria to adapt to acidic environments.

### Protonation of E136 Enhances Cytoplasmic Flexibility

To assess how E136 protonation influences channel flexibility, we analyzed the root mean square fluctuations (RMSF) of all residues, averaged over time, replicas, and protomers. The most pronounced differences were observed on the cytoplasmic side, where R20, and to a lesser extent E136, exhibited increased flexibility in the P-E154/E136 system compared to P-E154 (**Figure 5A**). This effect extended to the adjacent cytoplasmic loop, which also showed slightly enhanced mobility upon E136 protonation.

**Figure 5.**
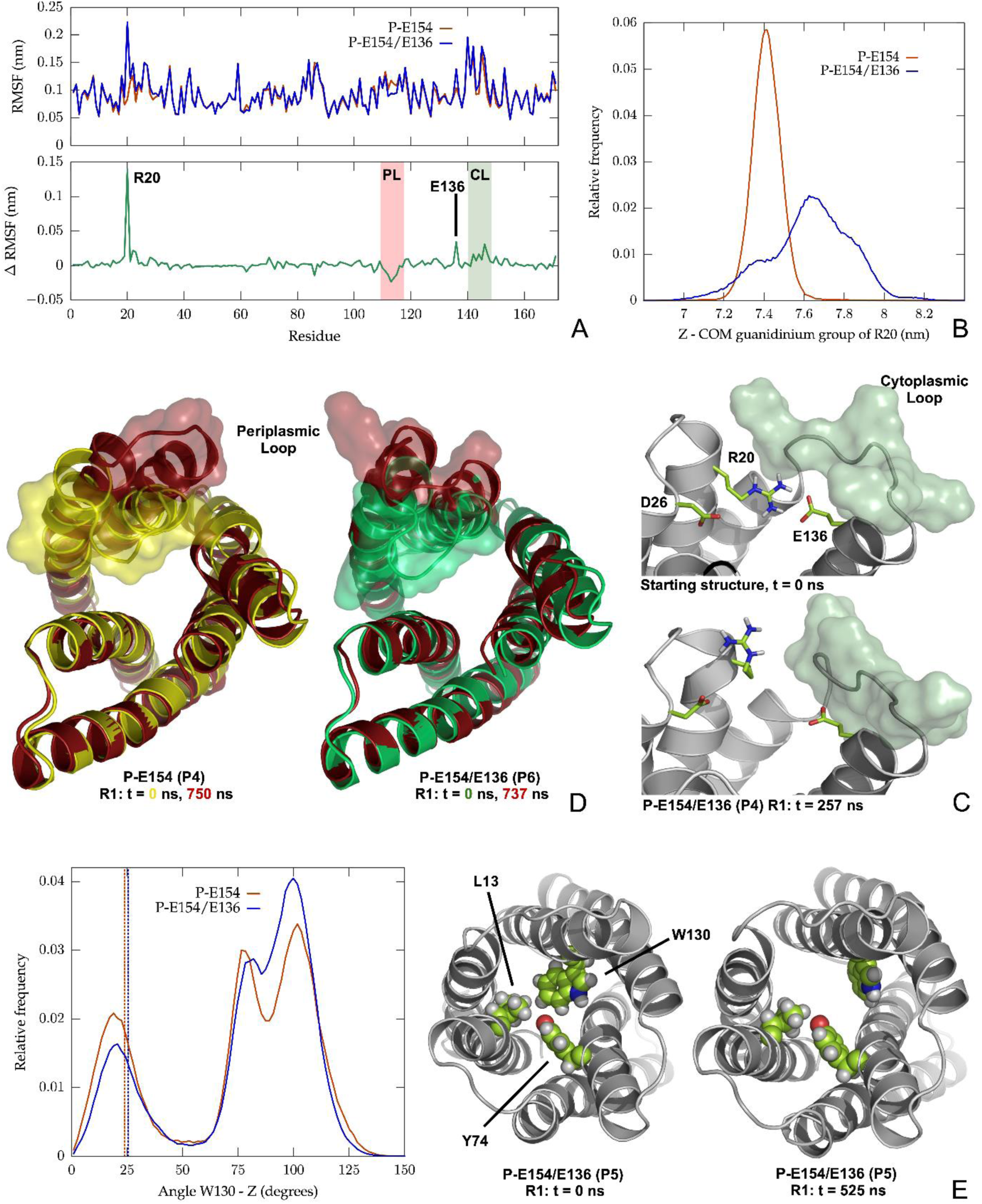
Conformational dynamics of SsUreI reveal pH-dependent gating and selectivity filter remodelling. **A.** Residue flexibility across protonation states. Top: Root mean square fluctuations (RMSF) of side chains, averaged over 100 ns windows (excluding the first 200 ns), replicas, and protomers for the P-E154 and P-E154/E136 systems. Bottom: Per-residue RMSF differences between the two systems, highlighting periplasmic (PL) and cytoplasmic (CL) loops. **B.** R20 mobility along the membrane normal. Distribution of the Z-coordinate (membrane normal) of the R20 guanidinium centre of mass across all replicas and protomers in both systems. Protonation of E136 increases R20’s positional variability, reflecting disruption of the E136–R20 salt bridge. **C.** Solvent-exposed conformation of R20. Representative snapshot from the P-E154/E136 simulation showing R20 in a solvent-exposed conformation, with no salt bridge to E136 or D26, compared to the starting model (top). The cytoplasmic loop is highlighted as a transparent surface. **D.** Periplasmic loop opening. Representative frames from P-E154 (left) and P-E154/E136 (right) simulations, superimposed on their starting structures, illustrating conformational rearrangements of the PL1 loop that widen the periplasmic entrance. **E.** W130 orientation. Left: Distribution of the angle between the indole plane of W130 and the membrane normal for both systems, with starting values indicated (dotted lines). Centre: Starting structure of a monomer with the L13–Y74–W130 triad shown as spheres. Right: Snapshot from P-E154/E136 illustrating a W130 rotation toward 90°, a conformation potentially favouring urea passage.

The enhanced flexibility of R20 resulted in large displacements of its guanidinium group, which oscillated by >1.4 nm along the membrane normal in the P-E154/E136 system (**Figure 5B**). In contrast, in the P-E154 simulations, the guanidinium group remained confined within a narrow range (∼few angstroms) due to the persistent E136–R20 salt bridge. Protonation of E136 also shifted the average position of R20’s guanidinium group toward the solvent-exposed region of the cytoplasmic vestibule (**Figure 5C**), though it still occasionally sampled more buried positions within the channel lumen. The >1.4 nm displacement of R20’s guanidinium group in the P-E154/E136 system suggests that E136 protonation may destabilize the cytoplasmic gate, enabling urea permeation

### Periplasmic Loop Rearrangements Suggest a Constitutively Open State

On the periplasmic side, we observed conformational rearrangements of the PL1 loop in a limited subset of monomers: 4 out of 24 monomers in the P-E154 simulations (6 protomers × 4 replicas); 1 out of 24 monomers in the P-E154/E136 system. These rearrangements involved a displacement of the loop backbone, widening the periplasmic entrance of the channel (**Figure 5D**). Strikingly, the PL1 loop in *Ss*UreI corresponds to PL1 in *Hp*UreI, which undergoes a major shift between its closed and open cryo-EM states^11^ (**Figure S9**). While the low frequency of these events likely reflects insufficient sampling of rare transitions in microsecond-scale simulations, their occurrence in both protonation states suggests that periplasmic loop opening is pH-independent. This implies that the physiologically relevant conformation of *Ss*UreI’s periplasmic side is constitutively open. Future simulations, starting from open-like conformations and extending sampling times, will be required to validate this hypothesis.

### W130 Dynamics Support a Role in Urea Selectivity and Permeation

Focusing on the internal constriction of the channel, we monitored the orientation of aromatic residues (W123, W126, Y74, W130) lining the selectivity filter. The angle between the indole/aromatic plane normal and the membrane normal (Z-axis) was used to assess permeability-favorable conformations (90° = perpendicular to membrane, favoring urea passage; deviations towards more parallel orientations with the membrane plane, disfavoring passage).

While W123, W126, and Y74 showed no systematic differences between protonation states (**Extended Data Figure 6**), W130 exhibited marked variability (**Figure 5E**). In the P-E154/E136 system, W130 sampled a larger fraction of 90° conformations, suggesting that E136 protonation may enhance urea permeation by dynamically remodeling the selectivity filter. Together, these results suggest a pH-dependent cytoplasmic gating (via E136/R20) and a constitutively open periplasmic gate, with W130 acting as a dynamic selectivity filter.

## Discussion

Our study reveals that *Ss*UreI functions as a pH-gated urea and ammonia channel, challenging the prevailing assumption that *Ss*UreI, and potentially Gram-positive UreI channels, operate independently of pH^12^. Using a combination of yeast complementation assays, in vitro permeability measurements, and all-atom MD simulations, we demonstrate that *Ss*UreI’s unitary urea permeability (47 ± 6 × 10⁻¹⁸ cm³/s) is three orders of magnitude lower than the unitary water permeability of water channels like aquaporins^15,19,26^, reflecting its specialized role in selective urea transport. This distinction underscores the diverse selectivity mechanisms underlying transport requirements for urea versus water in biological membranes. Furthermore, it establishes a quantitative benchmark for comparing urea permeability across UreI homologs and other urea transporters, including UTs^32^, aquaglyceroporins^33^, and ABC-type systems^34^.

A key finding is that pH-dependent gating in *Ss*UreI is mediated by the cytoplasmic residue E136, which acts as a conserved pH sensor across bacterial UreI channels. This contrasts with the prevailing model for *Hp*UreI, where periplasmic histidines were proposed as the sole pH sensors^10,11^. Our mutagenesis studies in *Ss*UreI, *H. hepaticus* UreI, and *H. pylori* UreI demonstrate that E136 (and its homologs E135/E159) is critical for pH gating, while R20 (or T170 in *H. pylori*) stabilizes the interaction. This suggests a dual-sensor mechanism in Gram-negative bacteria, where periplasmic histidines may complement cytoplasmic E136 to sense pH across the periplasmic space, a feature absent in Gram-positive bacteria like *S. salivarius*, which lack extensive periplasmic loops.

MD simulations provided molecular insights into the gating mechanism, showing that protonation of E136 disrupts its salt bridge with R20, increasing conformational flexibility at the cytoplasmic exit and facilitating urea release. Additionally, E136 protonation enhances the sampling of W130 conformations compatible with urea permeation, supporting a model where cytoplasmic pH sensing regulates channel activity. Notably, W130 is part of a conserved aromatic triad (L13/Y74/W130 in *Ss*UreI; L13/Y88/W153 in *Hp*UreI) that lines the selectivity filter^11^, and its dynamics are consistent with transient pore opening, as previously proposed for HpUreI^35^. The observation that periplasmic loop rearrangements occur independently of E136 protonation further suggests that the periplasmic gate remains constitutively open, while cytoplasmic gating is pH-regulated.

These findings, enabled by the use of *Ss*UreI as a simplified model system lacking periplasmic histidines, reveal a conserved yet adaptable mechanism for pH-dependent transport in bacterial UreI channels. In Gram-positive bacteria, E136 functions as the sole pH sensor, whereas Gram-negative bacteria may employ a dual-sensor model involving both cytoplasmic (E136) and periplasmic (histidine) residues. This adaptation likely reflects the need to sense and respond to periplasmic acidity in more complex bacterial cell envelopes. Given the critical role of UreI channels in acid resistance and virulence, targeting the E136–R20 interaction could provide a novel therapeutic strategy against antibiotic-resistant pathogens like *H. pylori*. Moreover, our results underscore the physiological relevance of pH-dependent urea transport in *S. salivarius*, complementing pH-regulated urease activity to maintain cytoplasmic pH homeostasis^36^ (**Figure 6**).

**Figure 6.**
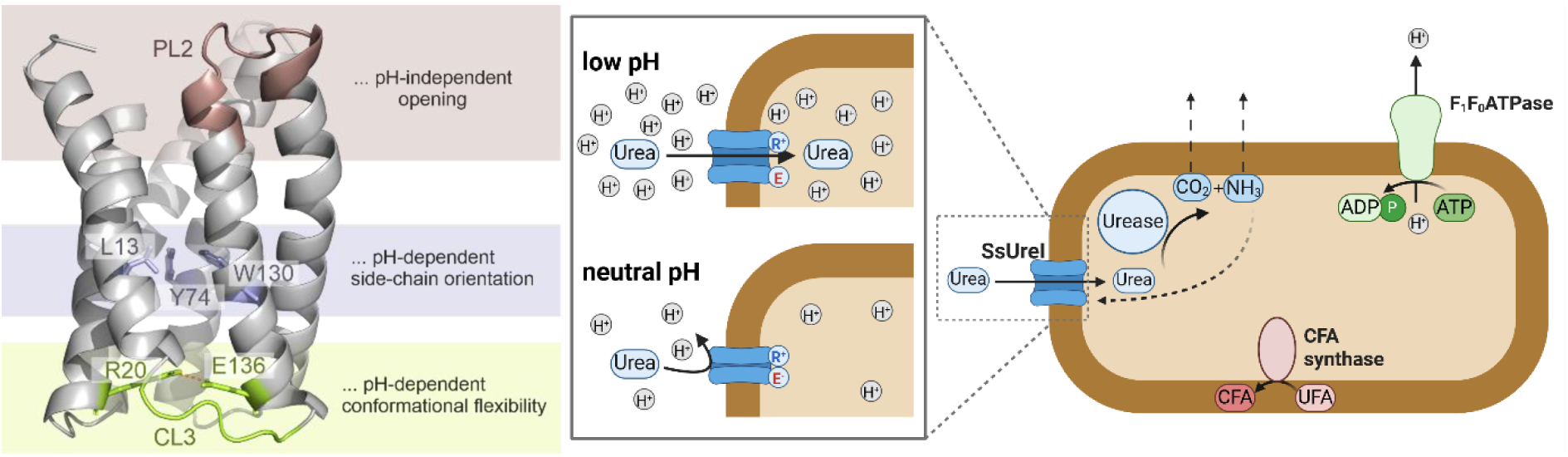
Molecular mechanism of pH-dependent urea uptake in Streptococcus salivarius acid acclimation. Structural basis of pH-dependent gating in SsUreI. Protonation of E136 disrupts the E136–R20 salt bridge, increasing the conformational flexibility of the cytoplasmic loop 3 (CL3) and W130 in the selectivity filter. This remodeling facilitates urea permeability through SsUreI, enabling rapid urea uptake under acidic conditions. Physiological role of SsUreI in acid resistance. SsUreI senses cytoplasmic pH in S. salivarius, supplying urea to the cytoplasm when pH declines. Urea uptake supports urease activity, maintaining intracellular pH, ensuring bacterial survival and growth in acidic environments, and stabilizing oral pH.

## Materials and Methods

### In vitro characterization

#### Plasmid Construction

The codon-optimized *SsUreI* gene (UniProt: J7T615) was cloned into a pET-26b vector (kanamycin resistance) with an N-terminal pelB-His₆-MBP tag, cleavable by HRV 3C protease, for affinity purification.

#### Transformation and Expression

The plasmid was transformed into chemically competent *E. coli* BL21(DE3) via heat shock. Transformants were selected on LB agar plates containing 50 µg/ml kanamycin overnight at 37°C. A single colony was inoculated into 50 ml 2×YT medium (200 µg/ml kanamycin) and grown overnight at 37°C (140 rpm). The preculture was diluted 1:20 into 1 L 2×YT medium (100 µg/ml kanamycin) and incubated at 37°C (140 rpm) until OD₆₀₀ = 0.6–0.8. Protein expression was induced with 1 mM IPTG, and cultures were incubated overnight at 28°C (140 rpm). Cells were harvested by centrifugation (6,500 × *g*, 20 min, 4°C), flash-frozen, and stored at −80°C.

#### Membrane Preparation

Thawed cell pellets were resuspended in lysis buffer (50 mM Na₂HPO₄, 150 mM NaCl, 1 mM MgCl₂, 1:500 DNase I, 1× cOmplete EDTA-free protease inhibitor, pH 7.4; 40 ml per 10 grams of pellet) and homogenized using an EmulsiFlex-C5 (Avestin). Unbroken cells and debris were removed by centrifugation (15,000 × *g*, 30 min, 4°C), and membranes were pelleted by ultracentrifugation (29,000 × *g*, 1.5 h, 4°C). Membranes were resuspended in resuspension buffer (50 mM Na₂HPO₄, 150 mM NaCl, pH 7.4; 2 ml per gram of cells) using a Dounce homogenizer, snap-frozen, and stored at −80°C.

#### Purification

The purification of *Ss*UreI followed a protocol similar to that employed for *Hp*UreI^37^. In summary, the membranes were thawed and solubilized in 1% (w/v) LDAO (Sigma-Aldrich) for 1 h at 4°C with stirring. Insoluble material was removed by ultracentrifugation (29,000 × *g*, 30 min, 4°C). Solubilized protein was bound to 1 ml Ni-NTA beads (Cube Biotech) pre-equilibrated in equilibration buffer (50 mM Na₂HPO₄, 150 mM NaCl, 0.4% LDAO, pH 7.4) and incubated for 2.5 h at 4°C. Beads were washed with 200 ml wash buffer (equilibration buffer + 10 mM imidazole) and 200 ml high-salt wash buffer (50 mM Na₂HPO₄, 750 mM NaCl, 0.4% LDAO, pH 7.4). Endogenous cysteines in PL1 were labeled with 50 µM Alexa Fluor 647 maleimide (5 ml, 1 h, RT) in labeling buffer (equilibration buffer + 400 µM TCEP). Excess dye was removed with 150 ml equilibration buffer, and protein was eluted in five 1-ml fractions of elution buffer (equilibration buffer + 250 mM imidazole). The pelB-His₆-MBP tag was cleaved overnight at 4°C using 40 U HRV 3C protease (Thermo Fisher Scientific).

#### Size-Exclusion Chromatography (SEC) and Quantification

Eluted protein was further purified by SEC (Superdex 200 Increase 10/300 GL, Bio-Rad) in equilibration buffer. Peak fractions (11–12.5 ml) were pooled, concentrated (100 kDa MWCO), and quantified using a detergent-compatible Bradford assay (Thermo Scientific). Purified *Ss*UreI was flash-frozen in 12.5% (v/v) glycerol and stored at −80°C.

#### Western Blot

Uncut *Ss*UreI samples were resolved by 15% SDS-PAGE (140 V, 1 h) and transferred to nitrocellulose (15 V, 70 min). Blots were blocked (5% skim milk powder in TBS, pH 7.4), probed with anti-His HRP antibody (1:5,000, Miltenyi Biotec) overnight at 4°C, and developed using ECL substrate (Abcam). Luminescence was detected using a Gel Imaging System (Bio-Rad). A representative blot is shown in Supplementary **Figure S3**.

#### Reconstitution into Proteoliposomes

Polar *E. coli* lipid extract (Avanti) was dried under nitrogen, resuspended in rehydration buffer (100 mM NaCl, 5 mM MOPS, 20 mM Na-acetate, pH 4.0–7.0) to 10 mg/ml, and vortexed to form multilamellar vesicles. Lipids were diluted to 5 mg/ml with detergent buffer (rehydration buffer + 7 mM DM (Anatrace)), and 0.5–1 mg/ml *Ss*UreI was added (1 h, RT). Detergent was removed by stepwise addition of Bio-Beads SM-2 (600, 750, and 1,000 µl). Proteoliposomes were extruded 21× through 100 nm polycarbonate filters (Avanti Mini-Extruder). Control vesicles (no protein) were prepared identically. Vesicle size and monodispersity were confirmed by DLS (DELSA Nano HC, Beckman Coulter)*^19,38^*.

### Quantitative Permeability Estimation

#### Water Permeability

Water permeability *P*_*f*_ was measured using stopped-flow light scattering (µSFM, Bio-Logic) at 5°C. PLUVs and control vesicles were mixed 1:1 (v/v) with hyperosmotic solution (300 mM sucrose, 100 mM NaCl, 20 mM MOPS, pH 7.4). The 90° scattered light intensity (λ = 546 nm) was recorded to track vesicle volume changes*^14-18,38^*. *P*_*f*_ was calculated using the analytical solution*^15,19,20^*:

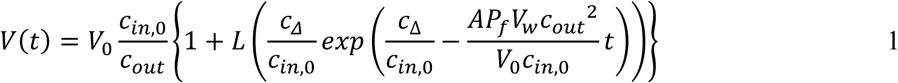

where *V*(*t*) = vesicle volume at time *t*, *V*_0_ = initial vesicle volume, *c*_*in*,0_ = initial internal osmolarity, *c*_*out*_ = external osmolarity, *c*_Δ_ = *c*_*out*_ − *c*_*in*,0_ (osmolyte gradient), *A* = vesicle surface area, *V*_*w*_ = molar volume of water (18 cm³/mol), and *L*(*x*) = Lambert function (*x*) · *e*^*L*(*x*)^ = *x*.

#### Urea Permeability

Urea permeability was assessed under isosmotic conditions at 5°C using the stopped-flow apparatus. Vesicles were pre-equilibrated with 300 mM urea (1:1 (v/v) dilution of 600 mM urea in rehydration buffer, overnight at 4°C). The mixing buffer was adjusted to match vesicle osmolarity using sucrose (verified with a Semi-Micro Osmometer, Knauer). Urea efflux (followed by water efflux) was monitored via 90° light scattering (λ = 546 nm).

Solute permeability *P*_*s*_ was derived from the time constant (*τ*) of an exponential fit to the scattering data^19,39^:

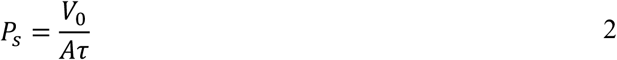

Unitary urea permeability *p*_*s*_ was determined by fluorescence correlation spectroscopy (FCS)^14,15,17^, accounting for contributions from the lipid bilayer *P*_*s*,*m*_ and incorporated channels *P*_*s*,*c*_:

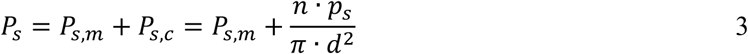

where *n* = the number of *Ss*UreI monomers per PLUV, quantified via AF647 fluorescence before/after micellization with 2% octylglucoside (OG) + 2% SDS), and *d* = vesicle diameter, respectively.

### Yeast Complementation Assays

#### Yeast Strains and Plasmids

For urea transport assays, we used the *S. cerevisiae* deletion strain YNVW1 (Δdur3), which lacks the endogenous urea transporter *DUR3*. Ammonia transport was assessed in the Sc18-Δmep1-3 strain, deficient in the endogenous ammonia transporters *MEP1-3*. Freeze survival assays employed the Y20000 strain (Euroscarf), which lacks functional endogenous *AYP*.

The genes encoding *Ss*UreI (UniProt: J7T615), *Hp*UreI (UniProt: P56874), *Hh*UreI (UniProt: G5EB92) and *human* AQP1 (UniProt: P29972; with a C-terminal 10×His tag) were codon-optimized for yeast expression and cloned into the galactose-inducible pYES2 vector. The *Ss*UreIcl construct included an N-terminal GPGSGS linker (residual sequence after HRV protease cleavage). Point mutations in *Ss*UreI were introduced via site-directed mutagenesis (Q5 High-Fidelity 2× Master Mix, NEB) using primers listed in Supplementary **Table S1** and verified by Sanger sequencing (Eurofins).

Plasmids were transformed into competent yeast cells using the Yeast Transformation Kit (Sigma-Aldrich), following a lithium acetate-based protocol^55^. Briefly, cells were incubated with 100 mM lithium acetate, mixed with carrier DNA and PEG buffer, and heat-shocked at 42°C for 15 min. Positive transformants were selected on DOB-ura plates.

#### Urea Growth Assay

Single colonies were inoculated overnight in 5 ml DOB-ura at 30°C with shaking (**Error! Reference source not found.A**). Cultures were diluted to 3.0 × 10⁶ cells/ml, washed with dH₂O to remove residual glucose, and induced overnight in galactose media (2% galactose, Yeast Nitrogen Base without ammonium sulfate or amino acids, 1 mM arginine, CSM-ura) at 30°C. Induced cells were adjusted to 4.0 × 10⁶ cells/ml in urea pH media (2% galactose, Yeast Nitrogen Base without ammonium sulfate or amino acids, 100 mM succinate-Bis-Tris-MOPS, 2 mM urea, pH 4.0–7.0) or arginine control media (1 mM arginine, pH 6.0). Cultures were incubated at 30°C for 48 h, with media replacement every 24 h. Cell density was measured in duplicate every 2.5 h using a CellDrop FL cell counter (DeNovix). The workflow is summarized in **Error! Reference source not found.B**.

#### Ammonia Growth Assay

The ammonia assay followed the same protocol as the urea assay but used ammonia pH media (2% galactose, Yeast Nitrogen Base without ammonium sulfate or amino acids, 100 mM succinate-Tris Base, 2 mM NH₄Cl, pH 4.0–7.0) or arginine control media (1 mM arginine, pH 6.0).

#### pK_a_ Estimation

Cell counts for each culture were normalized to the arginine control (pH 6.0). pH-dependency data (**Figures 2C, 3C, 4A+C+D**) were fitted to a sigmoidal Boltzmann function (OriginPro 2022):

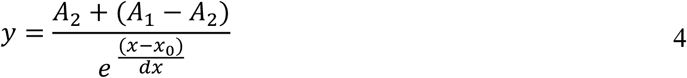

where *x*_0_represents the pKₐ (inflection point), and *A*_1_ and *A*_2_ denote the maximal and minimal permeabilities, respectively. Plots show the mean ± SEM from 3–18 independent assays. For *Ss*UreI point mutants, data were fitted linearly, and the intercepts are plotted in **Figure 3E**.

#### Water Freeze Assay

The workflow and principle for the freeze survival assay are depicted in Supplementary **Figures S1–S2**. Overnight cultures (DOB-ura, 30°C) were adjusted to 1.5 × 10⁷ cells/ml, washed, and induced for 24 h in galactose media. 7.5 × 10⁶ cells were resuspended in pH-buffered assay media (Yeast Nitrogen Base without ammonium sulfate or amino acids, 2% galactose, 0.5 M succinate-Bis-Tris-MOPS, pH 4.0–7.0). 40-µl aliquots were collected before and after a double freeze-thaw cycle (1 min in liquid nitrogen, followed by 45 min thawing at room temperature).

Viability was assessed using a yeast viability kit (Logos BioSystem) with fluorescein-diacetate (FDA) and propidium iodide (PI). FDA stains viable cells green (metabolized by esterases), while PI stains damaged cells red. After 10-minute incubation with a fluorescence enhancer in the dark, stained samples were analyzed using the CellDrop FL cell counter (DeNovix). Normalized viability was calculated as:

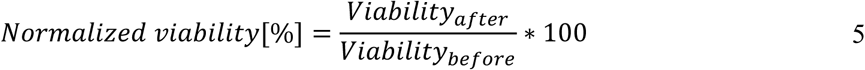

Relative viability was determined by subtracting the normalized viability of the empty vector control:

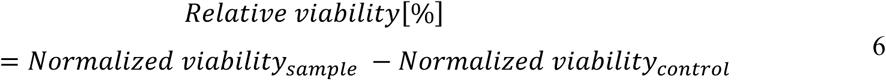

pH-dependency curves (**Extended Data Figure 1E**) were fitted sigmoidally using the Boltzmann function, while relative viability data (3–5 independent assays) were fitted linearly.

#### Molecular Dynamics Simulations

Molecular dynamics (MD) simulations were performed using the GROMACS software package with the CHARMM36m force field^40,41^. The *Ss*UreI hexamer structure, predicted using AlphaFold3^13^, was refined by manually adjusting inner lipids to fit the central plug using VMD tools^42^, following the protocol established by McNulty et al.^35^ for *Hp*UreI. The hexamer was embedded in a POPC bilayer, solvated in a 12.6 × 12.6 × 12.5 nm box with the TIP3P water model^43^ and 100 mM NaCl, resulting in systems of ∼205,000 atoms. The lipid arrangement included three POPC lipids in the periplasmic leaflet and nine POPC lipids in the cytoplasmic leaflet of the plug.

Three protonation states were simulated: standard protonation (SP), E154 protonated (P-E154; mimicking pH 7.5), and E154/E136 protonated (P-E154/E136; mimicking pH 5.0). For each system, four independent 1-µs replicas were run in the NpT ensemble with distinct starting velocities.

pKₐ calculations were performed using the PyPKa tool^44^, a Python module for Poisson-Boltzmann-based pKₐ predictions (DelPhi^45^). Detailed MD protocols and pKₐ calculation parameters are provided in Supplementary Methods.

#### Sequence- and Structure-Based Analysis

Homolog identification was performed using BLAST^46^ with input sequences for *S. salivarius* UreI (*Ss*UreI; UniProt: J7T615) and *H. pylori* UreI (*Hp*UreI; UniProt: Q09068). Multiple sequence alignments were generated using Clustal Omega^47^, and guide trees were visualized with iTOL^48^ (**Extended Data Figures 3** and **5**). Sequence conservation was analyzed using custom Python (3.13.0) scripts to create sequence conservation graphs (**Figure S6**) and sequence logos utilizing the logomaker package^49^ (**Figure 3D**).

## Acknowledgements

The financial support for this study is from the Austrian Science Fund (FWF) [P31074 and P33541] to AH. GP, MF, and DN acknowledge the CINECA award under the ISCRA initiative for the availability of high-performance computing resources and support. The authors thank Hartmut Lücke for generously providing the *Ss*UreIcl plasmid.

## Author contributions

AH conceived the project. CS, AS, and SS prepared the constructs. AS optimized and performed yeast complementation assays for urea and water. SS optimized and performed yeast complementation assays for ammonia. AS and SS prepared variant plasmids. SS performed urea and ammonia complementation assays with *Ss*UreI variants. AS analysed yeast complementation assays. XF and SP overexpressed, purified, and reconstituted *Ss*UreI into lipid vesicles and performed *in-vitro* experiments. TP performed sequence and structure analysis. AS, SP, and NGM are responsible for the artwork. DN and GP designed the molecular dynamics simulations. GP performed the molecular dynamics simulations and carried out the pK_a_ analysis of the resulting trajectories; MF performed the structural analysis on the MD trajectories. AS, AH, GP, MF, and DN wrote the manuscript; all the authors made manuscript revisions. DN conceived and supervised the theoretical part of the study.

## Data Availability Statement

Data required to support our conclusions are presented in the article, the Extended Data Items, and the Supplementary Information file. Data used to perform MD simulations and the experimental data substantiating the findings of this study will be deposited in Zenodo repositories upon publication.

## Extended Data Figures

**Extended Data Figure 1.**
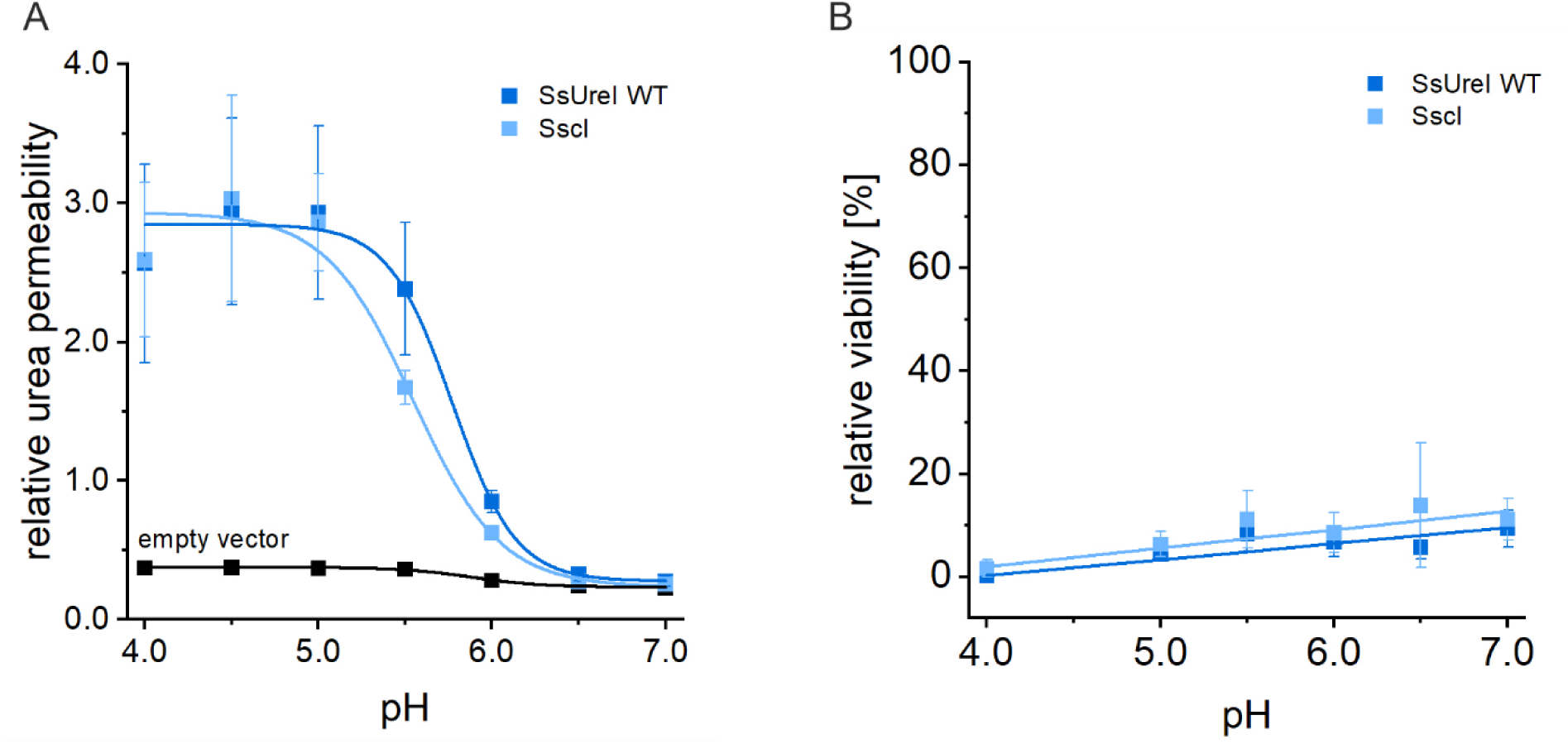
Functional comparison of wild-type SsUreI and the purification construct (SsUreIcl). **A.** Yeast complementation assays with urea as the sole nitrogen source demonstrate that SsUreIcl restores pH-dependent growth in S. cerevisiae Δdur3 cells, similar to wild-type SsUreI across pH 4.0–7.0. An empty vector serves as a negative control. **B.** Water freeze-thaw survival assays reveal that SsUreIcl (n=6) confers equivalent protection against freeze-induced damage (FDA/PI staining) as the wild-type channel (n=4). Data represent mean ± SEM.

**Extended Data Figure 2.**
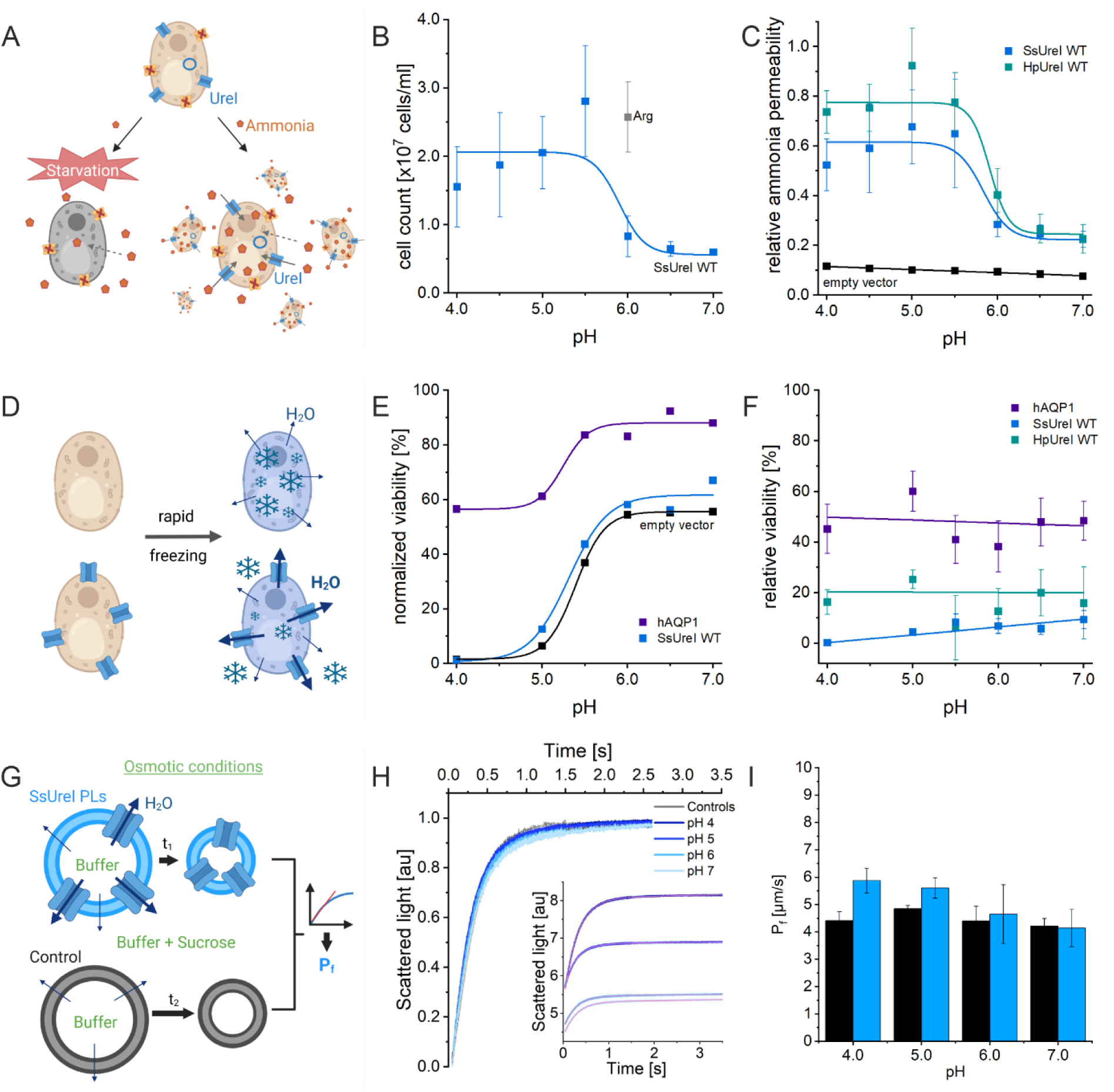
SsUreI selectively transports ammonia but exhibits negligible water permeability. **A.** Yeast growth assay for ammonia permeability. The ammonia uptake-deficient strain S. cerevisiae Δmep1-3 expressing SsUreI grows only if the channel facilitates ammonia transport. Higher ammonia permeability results in increased cell growth, while low membrane permeability (dotted arrows) leads to starvation. **B.** pH-dependent growth of SsUreI-expressing yeast. Cell counts were measured after 2 days of incubation in ammonia media at pH 4.0–7.0 or arginine media at pH 6.0 (internal standard, gray). Data represent exemplary average (three times as duplicates) cell counts ± SD for SsUreI WT in ammonia (blue) and arginine (gray) media. **C.** Normalized ammonia permeability of SsUreI and HpUreI. pH-dependent relative ammonia permeability for SsUreI (n=3) and HpUreI (n=7). Unlike urea permeability (Figure 2C), ammonia permeability is reduced but not abolished at neutral pH. **D.** Freeze-survival assay for water permeability. Cells with higher water permeability exhibit enhanced freeze survival due to reduced ice crystal formation. Viability is normalized to pre-freezing levels to account for baseline cell death. **E.** Exemplary freeze-survival raw data. Viability of yeast expressing empty vector (negative control), SsUreI WT, or hAQP1 (positive control) was determined using FDA (live cells, green) and PI (dead cells, red) as a technical duplicate. Viability is calculated as the percentage of live cells relative to total stained cells. **F.** pH-dependent freeze survival indicates minimal water permeability. Relative freeze survival of SsUreI and HpUreI is marginal compared to hAQP1, suggesting negligible water transport through UreI channels. Data represent averaged relative survival rates ± SEM from independent assays. **G.** Vesicle shrinkage assay for water permeability. Vesicles exposed to an osmotic gradient undergo water efflux, with the shrinkage rate correlating to water permeability *P*_*f*_. **H.** Exemplary vesicle shrinkage data. Scattered light signals from SsUreI PLUVs at pH 4.0, 5.0, 6.0, and 7.0 were fitted (red line) to calculate *P*_*f*_. (Inset) Raw data show minimal shrinkage, indicating low water permeability. **I.** Quantified water permeability of SsUreI PLUVs. *P*_*f*_ through SsUreI PLUVs (blue) is only marginally higher than control vesicles (black). Data represent mean ± SD from three independent experiments.

**Extended Data Figure 3.**
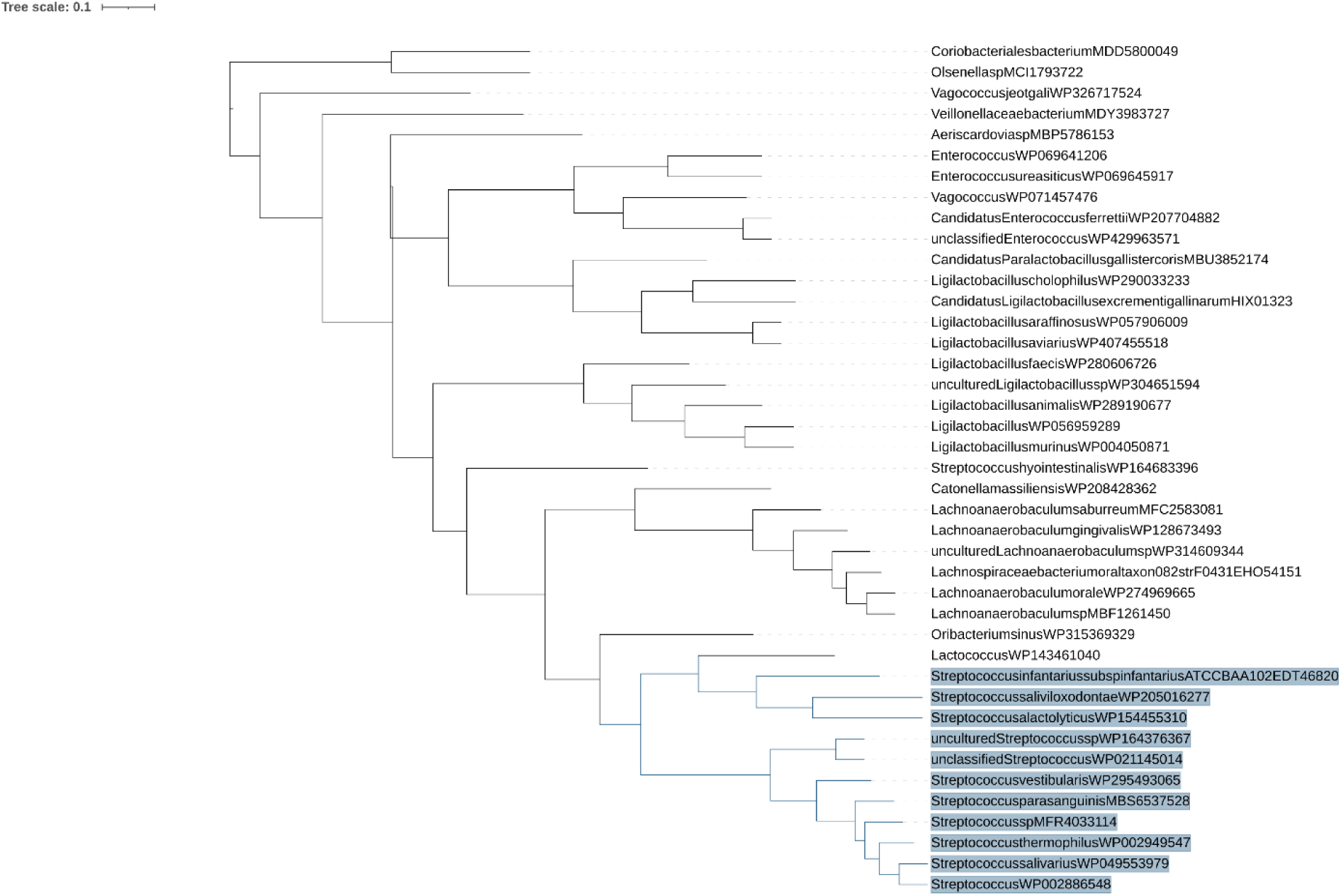
Guide tree of SsUreI homologs. A sequence-based guide tree was generated from a multiple sequence alignment of SsUreI and its closest homologs. The Streptococcus family (highlighted in blue) forms a distinct cluster, indicating a shared evolutionary origin of urea channels within this genus. Branch lengths are proportional to sequence divergence.

**Extended Data Figure 4.**
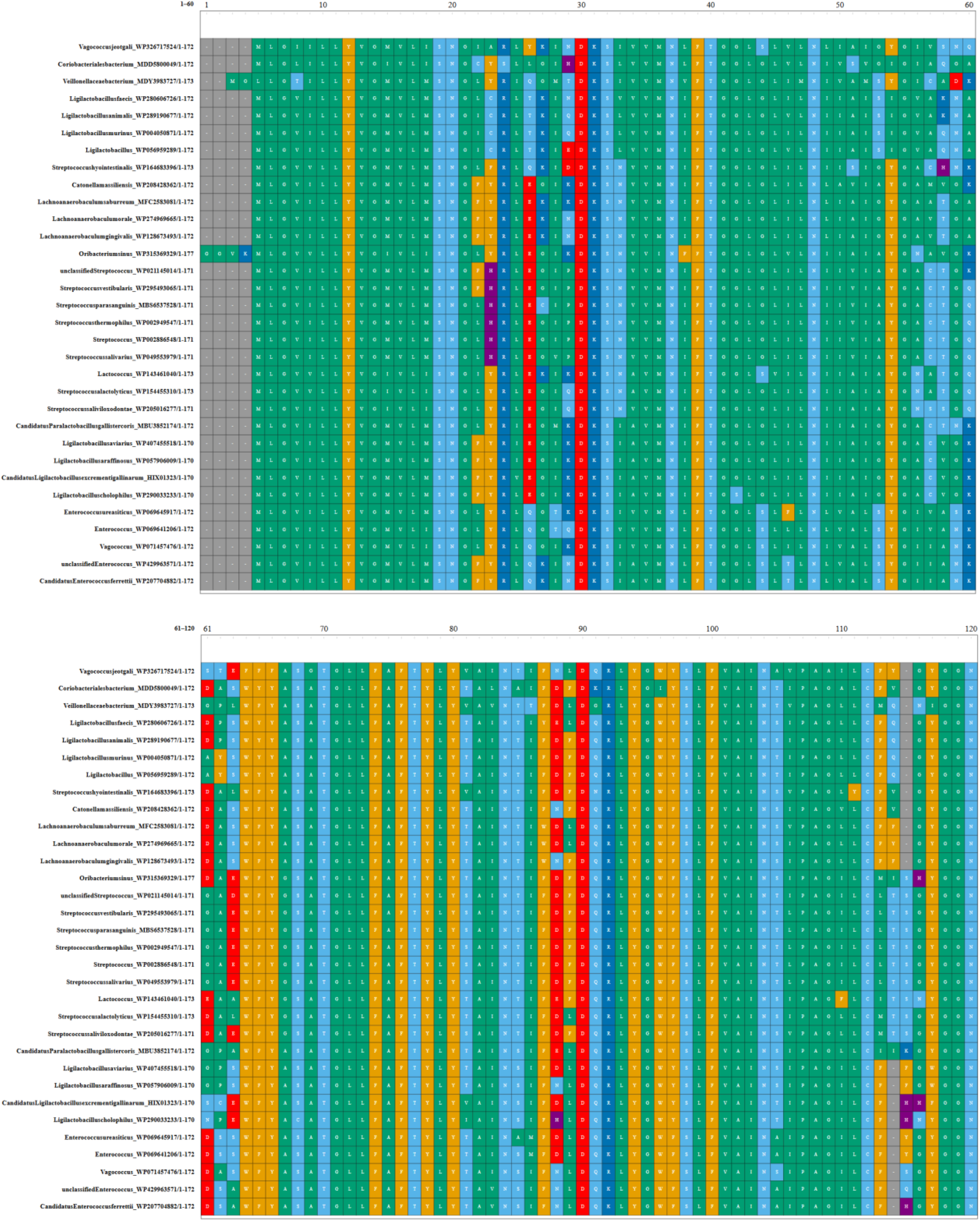
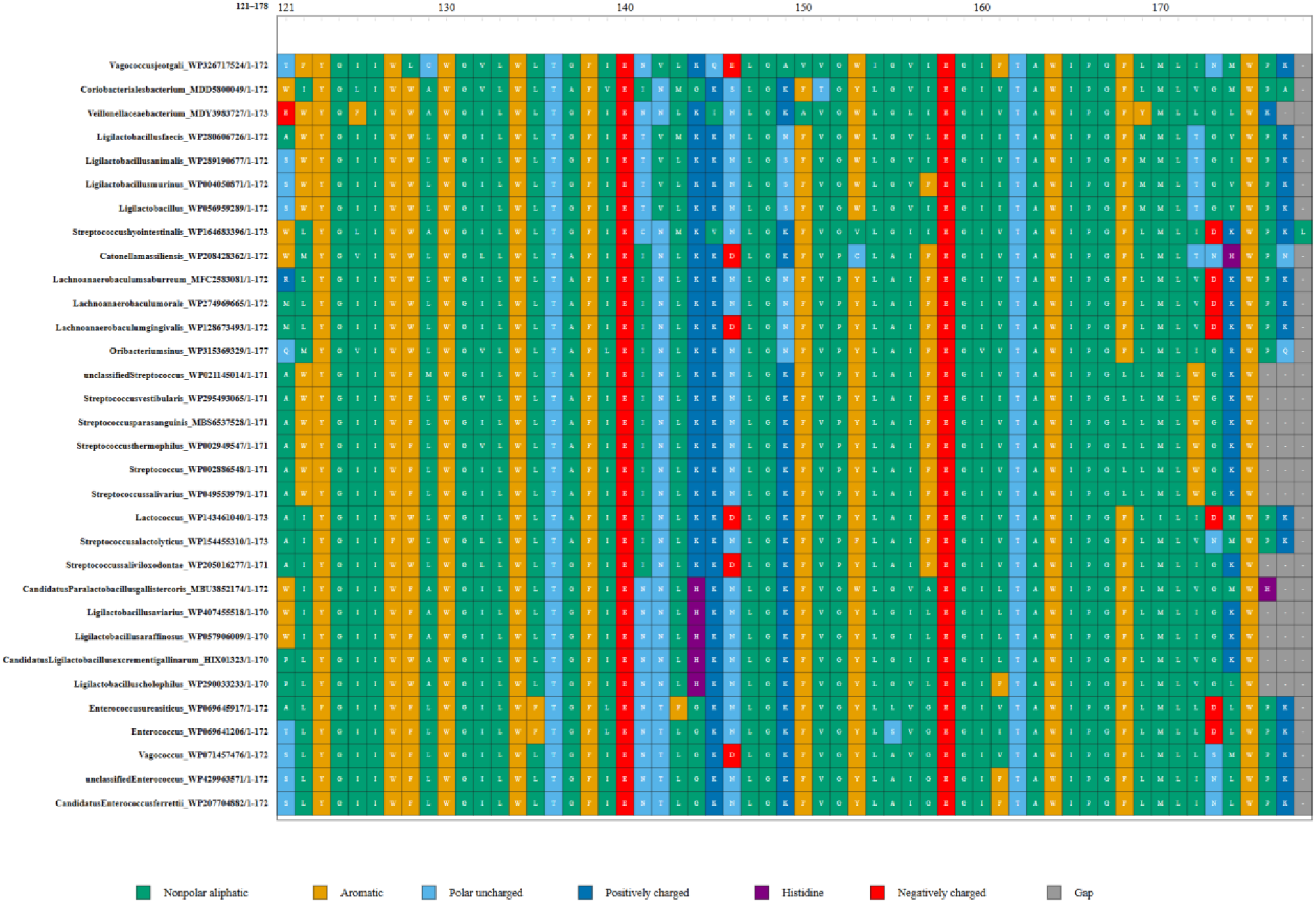
Multiple sequence alignment of SsUreI homologs. A multiple sequence alignment (MSA) highlights conserved regions among SsUreI and its homologs, with particular conservation of charged residues (e.g., E136 in SsUreI). Due to variable N-terminal lengths, the equivalent of SsUreI’s E136 appears at position 140. Accession numbers for each sequence (corresponding to **Extended Data** Figure 3) are listed on the left, with residues color-coded depending on their chemical property (see legend below).

**Extended Data Figure 5.**
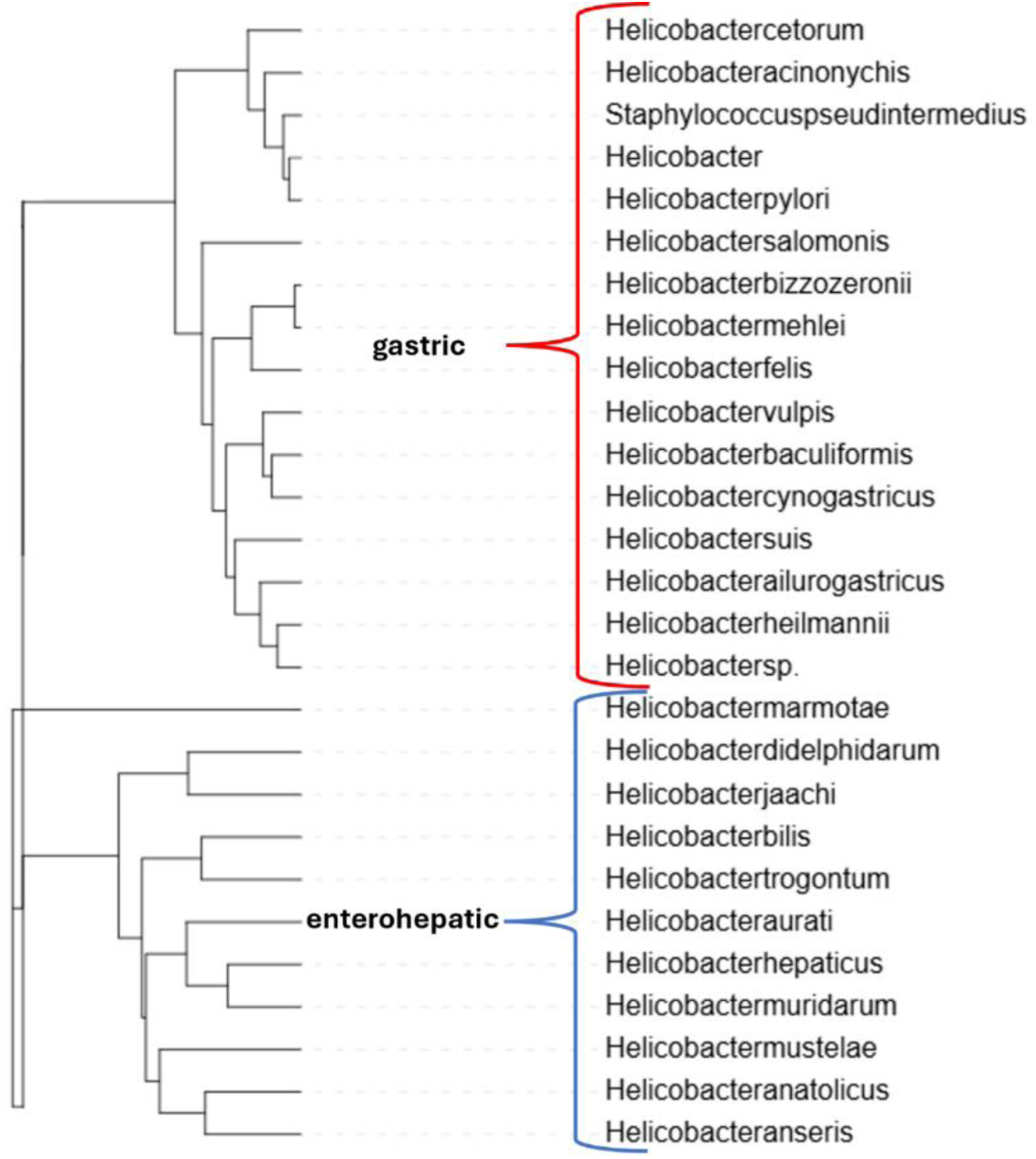
Phylogenetic clustering of UreI homologs from Helicobacter species. A guide tree of UreI homologs reveals two distinct clusters corresponding to: Gastric Helicobacter (e.g., H. pylori), characterized by long periplasmic loops; Enterohepatic Helicobacter (e.g., H. hepaticus) characterized by short periplasmic loops^50^. This division correlates with ecological niche specialization and suggests divergent pH-sensing mechanisms between gastric (acidic stomach environment) and enterohepatic (neutral/bile-rich environment) pathogens.

**Extended Data Figure 6.**
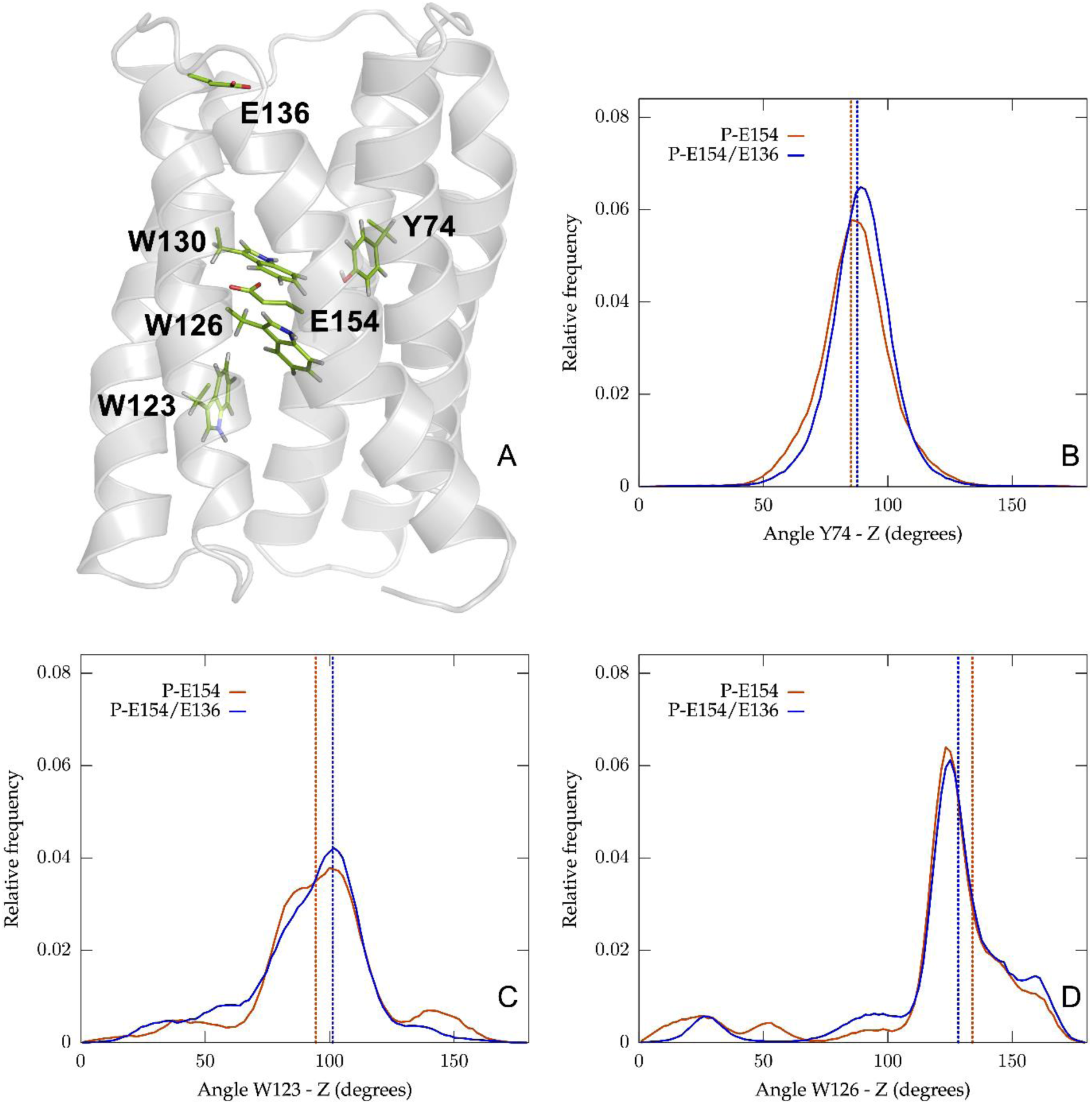
Conformational dynamics of key filter residues in SsUreI. **A.** Structural context: Side view of a single protomer (cartoon) with key residues (Y74, W123, W126, W130, E154, E136) shown in stick representation. **B–D.** Residue orientation analysis: W123 (B), W126 (C), Y74 (D). Distributions of the angle between the ring plane normal (indole for W123/W126, phenolic for Y74) and the membrane normal are shown for both protonation states (P-E154: red lines; P-E154/E136: blue lines). Starting angles are indicated by dotted lines, illustrating conformational shifts during simulations.

**Extended Data Table 1.**
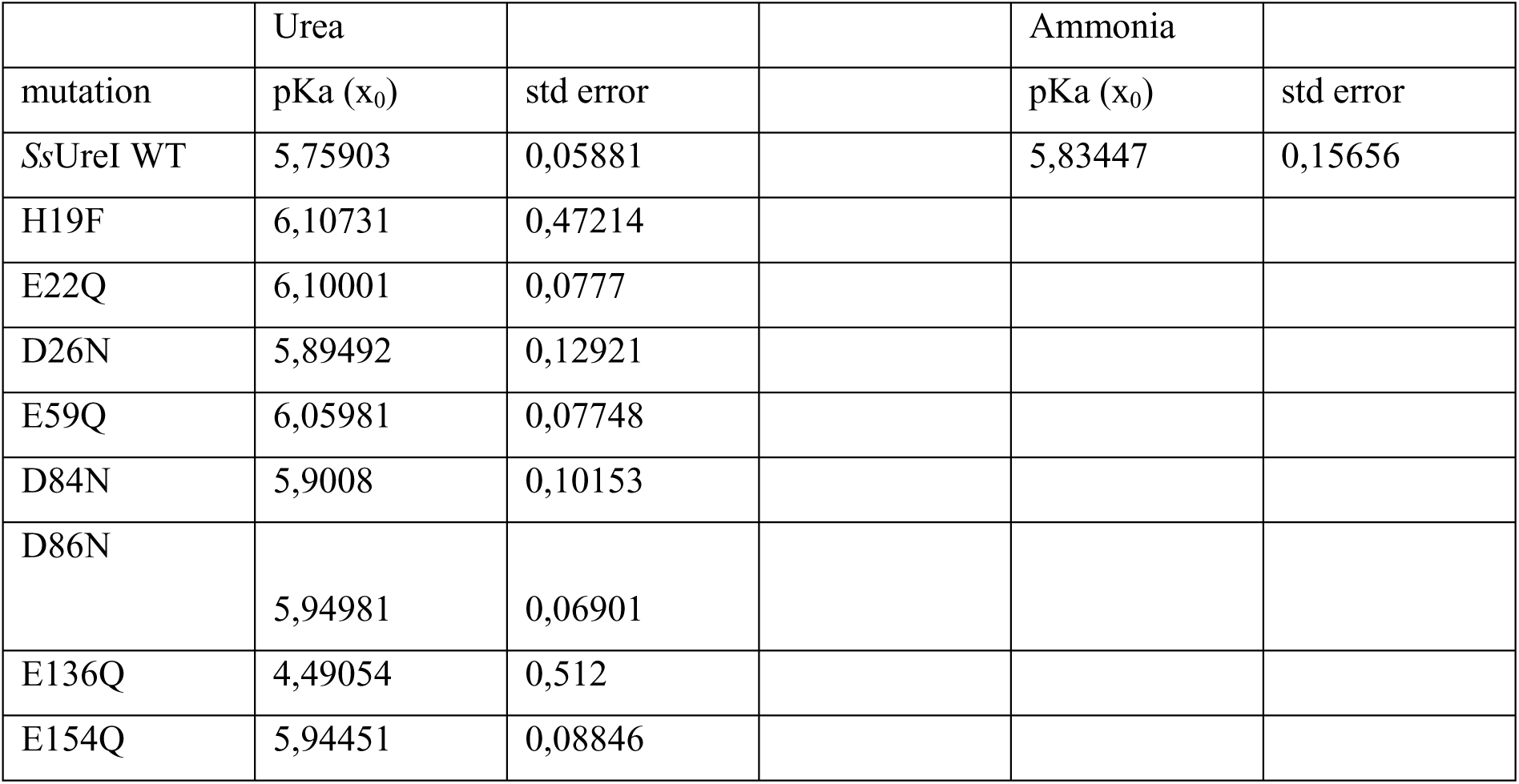
pKa values of SsUreI channel gating. pKₐ values correspond to the color-coded bar charts of relative urea and ammonia permeabilities for SsUreI variants (Figure 3A,B). Values were derived from sigmoidal fits of pH-dependency data (Boltzmann function).

**Extended Data Table 2.** Statistics of inter-residue contacts in SsUreI simulations. Inter-residue contacts were analyzed across three protonation states (SP, P-E154, P-E154/E136), averaging over four 1,000 ns trajectories and six protomers per system. Reported values include the average occupancy (fraction of frames where the inter-residue distance was ≤ 0.35 nm), the normalized formation/breaking ratio (Relative frequency of contact formation vs. disruption events), and the mean lifetime (Average duration of contact (ps)). These metrics reveal how protonation of E136 disrupts critical contacts (e.g., E136–R20, R88–E22), providing a structural basis for pH-dependent gating in SsUreI.

|  | occupancy | formation/breaking | lifetime (ps) |
| --- | --- | --- | --- |
| SP |  |  |  |
| Arg20-Asp26 | 0.026 | 0.005 | 44.548 |
| Arg20-Glu136 | 0.868 | 0.010 | 4217.122 |
| Glu22-Arg88 | 0.528 | 0.035 | 5162.718 |
| P-E154 |  |  |  |
| Arg20-Asp26 | 0.003 | 0.001 | 25.707 |
| Arg20-Glu136 | 0.984 | 0.042 | 67566.245 |
| Glu22-Arg88 | 0.573 | 0.022 | 547.622 |
| P-E154/E136 |  |  |  |
| Arg20-Asp26 | 0.159 | 0.042 | 36.437 |
| Arg20-Glu136 | 0.132 | 0.027 | 51.717 |
| Glu22-Arg88 | 0.339 | 0.031 | 251.671 |

